# A neural circuit arbitrates between perseverance and withdrawal in hungry *Drosophila*

**DOI:** 10.1101/259119

**Authors:** S. Sayin, J.-F. De Backer, M.E. Wosniack, L.P. Lewis, K.P. Siju, L.-M. Frisch, P. Schlegel, A. Edmondson-Stait, N. Sharifi, C.B. Fisher, S. Calle-Schuler, S. Lauritzen, D.D. Bock, M. Costa, G.S.X.E. Jefferis, J. Gjorgjieva, I.C. Grunwald Kadow

## Abstract

In pursuit of palatable food, hungry animals mobilize significant energy resources and overcome obstacles, exhaustion and fear. Their perseverance depends on metabolic state, internal motivation and the expected benefit. Sustained commitment to a trying task is crucial, however, disengagement from one behavior to engage into another can be essential for optimal adaptation and survival. How neural circuits allow prioritizing perseverance over withdrawal based on the animal’s need is not understood. Using a single fly spherical treadmill, we show that hungry flies display increasing perseverance to track a food odor in the repeated absence of the predicted food reward. While this perseverance is mediated by a group of dopaminergic neurons, a subset of neurons expressing octopamine, the invertebrate counterpart of noradrenaline, provide reward feedback and counteract dopamine-motivated food seeking. Our data and modeling suggest that two important neuromodulators tally internal and external signals to coordinate motivation-dependent antagonistic behavioral drives: perseverance *vs.* change of behavior.

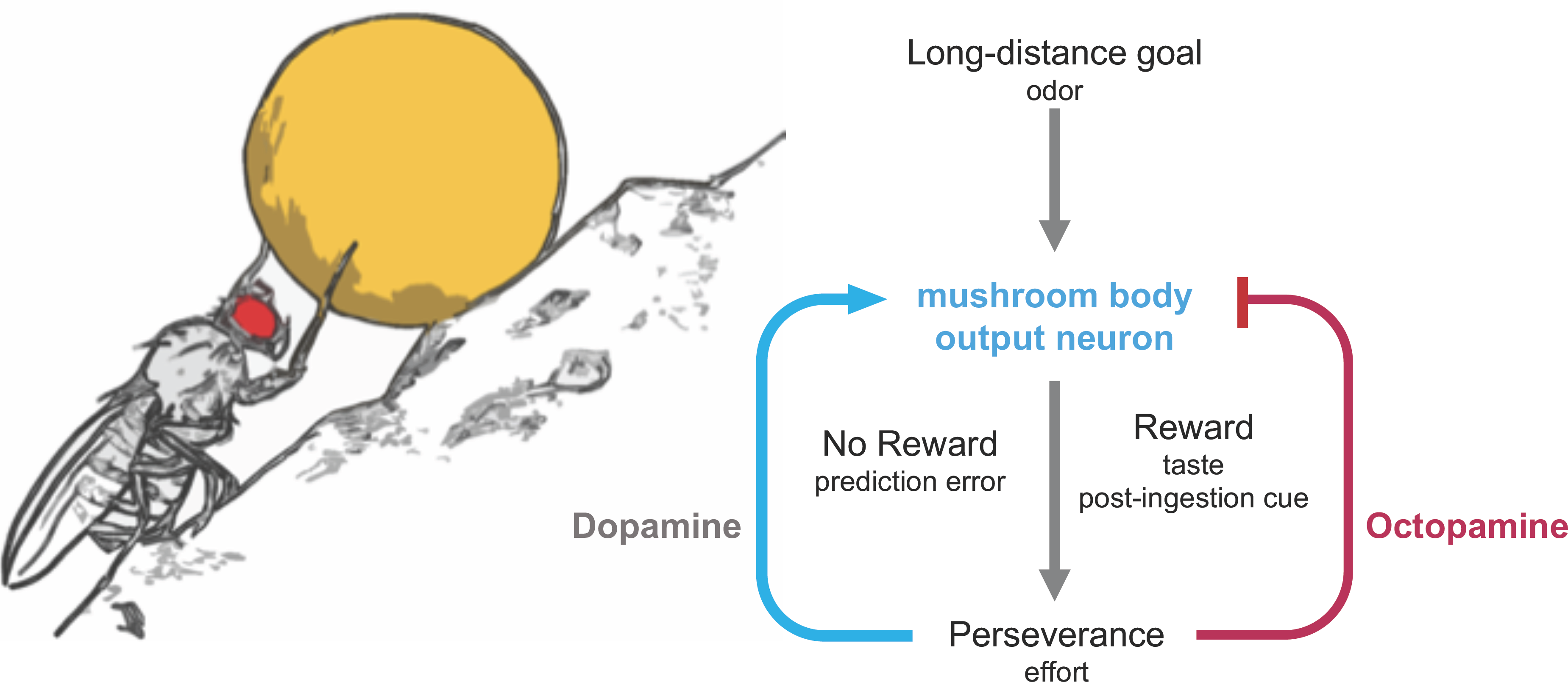

**Highlights:**

- Lack of reward stimulates perseverance, and not quitting.
- Dopaminergic neurons previously implicated in aversive learning promote perseverance.
- Sugar responsive octopaminergic neurons directly counteract perseverant odor tracking through a downstream inhibitory neuron.
- Computational modeling supports a simple neural circuit featuring antagonistic functions for dopamine and octopamine as tallies of expense and gain.

## Introduction

Flexibility is an important factor in an ever-influx environment, where scarcity and competition are the norm. Without perseverance to achieve its goals, an animal’s daily strive, to secure food, protect its offspring, and even to maintain its social status are in jeopardy. Therefore, sensory cues related to food or danger often translate into strong impulses in animals. However, these impulses must be strictly controlled to allow for coherent goal-directed behavior and to permit behavioral transitions when opportune. Inhibition of antagonistic behavioral drives at the cognitive and physiological level has been proposed as a major task of a nervous system (Bari and Robbins, 2013). Which sensory cues and ultimately which behavior is prioritized and wins, depends on the animal’s metabolic state, internal motivation and the expected reward. In mammals, norepinephrine (NE) released by a brain stem nucleus, the *locus coeruleus* (LC) has been implicated in controlling the balance between perseverance and action selection (Berridge and Waterhouse, 2003; Schwarz and Luo, 2015). Furthermore, NE neurons of the nucleus of the solitary tract (NST) innervate regions such as the central nucleus of the amygdala and hypothalamus. They receive taste information as well as input from the gastrointestinal tracts, lungs and heart to presumably mediate taste and control autonomic functions, respectively (Carleton et al., 2010). While earlier work treated these NE rich nuclei as uniform areas, recent efforts using the mouse aim at dissecting their diverse neural connections within the CNS and their intrinsic heterogeneity (Schwarz and Luo, 2015; Schwarz et al., 2015) in order to reveal their different functions.

The functional counterpart of norepinephrine in insects appears to be octopamine (OA). OA neurons (OANs) are organized in distinct clusters in the brain of *Drosophila melanogaster,* with one of the main clusters close to the primary olfactory input region, the antennal lobe (AL), and another ventral cluster within the subesophageal zone (SEZ), the main region of gustatory input (Busch and Tanimoto, 2010). This ventral cluster contains several types of OA neurons, some of which appear to receive information from ventral regions such as the SEZ and project axons to diverse higher brain regions, including the mushroom body (MB), in a cell type specific manner (Busch et al., 2009; Busch and Tanimoto, 2010). OA has been implicated in a number of behaviors including arousal, sleep, endurance, aggression, memory, and energy and nutrient homeostasis (Corrales-Carvajal et al., 2016; Crocker et al., 2010; Schwaerzel et al., 2003; Sujkowski et al., 2017; Watanabe et al., 2017; Yang et al., 2015). In addition, OA also modulates early sensory processing in adult flies and larvae (Berck et al., 2016; Strother et al., 2018; van Breugel et al., 2014). Outside of the brain, octopaminergic signaling is necessary at the neuromuscular junction and promotes increased locomotion under food deprivation (Koon et al., 2011). Flies lacking OA show reduced arousal upon starvation and accumulate fat reserves (Li et al., 2016; Shang et al., 2013; Yang et al., 2015; Zhang et al., 2013). Furthermore, OANs are important to form appetitive memories of odors (Burke et al., 2012; Perry and Barron, 2013). OA increases food intake and sensitizes sugar, and surprisingly also bitter taste neurons (LeDue et al., 2016; Wang et al., 2016), emphasizing the differential and even opposite roles octopaminergic neurons can play during state-dependent behavior. Although some of these studies identified specific neurons, the exact type of OAN involved in most contexts remained elusive.

Similar to NE and OA, dopamine is being studied in many aspects of behavioral adaptation, flexibility, and learning. In flies, the best-known role for dopaminergic neurons (DANs) stems from extensive research in olfactory learning and memory (Waddell, 2013). Here, different classes of DANs mediate negative or positive experience (Aso and Rubin, 2016; Burke et al., 2012; Liu et al., 2012; Riemensperger et al., 2005), novelty (Hattori et al., 2017) and forgetting (Berry et al., 2012). In addition, in mammals, DANs have been implicated as mediators of prediction errors in processes from reinforcement learning to economic decision-making (Schultz et al., 2017), but the mechanisms remain to be fully elucidated (Eshel et al., 2015; Watabe-Uchida et al., 2017). Recent data from *Drosophila* indicate that similar functions exist in flies (Felsenberg et al., 2017).

Like most animals, energy-deprived flies prioritize food seeking and feeding behavior over other behaviors such as mating, resting or hiding. To find food, flies can follow olfactory or visual cues over long distances. External gustatory cues provide information about the type of food the odor source presents. However, only internal nutrient levels will provide reliable feedback about the quality and quantity of a food source, and ultimately suppress food seeking behaviors (Dethier, 1976; Thoma et al., 2016; Yang et al., 2015). Therefore, food odor, the taste of food, and post-ingestion internal feedback signals induce sequential and antagonistic behaviors (Thoma et al., 2017) - perhaps in analogy to grooming behavior, which is organized in a highly sequential manner with hierarchical suppression of distinct motor programs (Seeds et al., 2014). Interestingly, these chemosensory and internal feedback systems appear to converge in the MB (Cohn et al., 2015; Krashes et al., 2009; Lewis et al., 2015). Which neurons and which signals combine external and internal cues to coordinate and suppress competing behavioral drives is not well understood.

Here, we analyzed the mechanistic relationship between perseverance in goal-seeking behavior and its suppression upon reaching the goal. Using a single fly spherical treadmill assay, we find that hungry flies increase their effort to track a food odor with every unrewarded trial rather than giving up. This perseverance depends on dopamine-mediated signals reinforcing behavioral performance, which are well captured in a model where DANs integrate effort and the lack of expected reward. Activation of sugar taste or internal sugar sensory neurons counteracts food seeking motivation briefly or lastingly, respectively, suggesting a sequential hierarchy between olfaction, gustation, and post-ingestion cues. At the neural circuit level, we pinpoint that a specific type of OAN, the VPM4 (ventral paired medial) neuron, mimics nutrient reward and suppresses MBON Y1pedc>α/β/MVP2-dependent odor tracking. Based on our experimental data and modeling, we propose that VPM4, perhaps in a somewhat similar role to the NST, acts as a bridge between internal state, taste information, and the higher brain, and thereby regulates the switch between food foraging and feeding.

## Results

### Flies persistently track attractive food odors in the absence of reward

To characterize perseverance in goal seeking behavior, we devised a spherical treadmill assay with a single tethered fly exposed to a low-speed frontal air or odor stimulus (Fig. 1A). Upon an initial period of 3 minutes of habituation, we recorded running speed, turns, and stops during a 20 s pre-stimulus, a 12 s stimulus, and a 20 s post-stimulus period (Fig. 1A, Fig. S1.1A). For each fly, the protocol was repeated 10 times with variable inter-stimulus intervals to minimize the likelihood that the fly could predict the exact onset of the stimulus. We used 3 ppm of vinegar odor as a highly attractive cue to analyze the behavior of a hungry fly (24 h starvation) when trying to reach the goal, the predicted food producing the odor (Fig. S1.1B). Flies ran on average at a speed of 7.3 mm/s during the pre-stimulus periods (Fig. 1B). During odor stimulation, flies sped up significantly and reached speeds of 12.4 mm/s on average (Fig. 1B). Upon cessation of the odor stimulus, flies showed a strong offset behavior with stopping before regaining an average speed of 6.8 mm/s (Fig. 1B). In addition to changing speed, the fly also suppressed turning to left and right and was headed straighter suggesting that it was indeed tracking the odorant (Fig. 1C). The loss of the olfactory cue at the end of the stimulation period led to a significant increase in turning behavior (Fig. 1C,D) suggesting that the animal was actively searching for the stimulus as previously observed for fly larvae (Gomez-Marin et al., 2011) and other animals such as dogs and even humans (Porter et al., 2007). Interestingly, this behavior evolved over the 10 trials. Although flies showed an initial acceleration at stimulus onset already during the first 3 trials, they did not track the odors at high speed for more than a fraction of the stimulus time (Fig. 1E, Fig. S1.1C). With increasing number of trials, the flies ran longer distances and more frequently for the entire stimulus time of 12 s (Fig. S1.1D,E). In addition, they ran faster with each trial and suppressed turning more efficiently (Fig. S1.1F-H). Nevertheless, the relative initial increase in speed from before to during odor stimulation was similar between trials 1 to 10 (R^2^=0.535, p<0.0001, n = 180 pairs). These data show that flies reliably track food odors by suppressing turning behavior and increasing speed. Furthermore, they suggest that in the absence of the expected food reward, flies do not give up on their goal, but instead become increasingly perseverant.

**Fig. 1:**
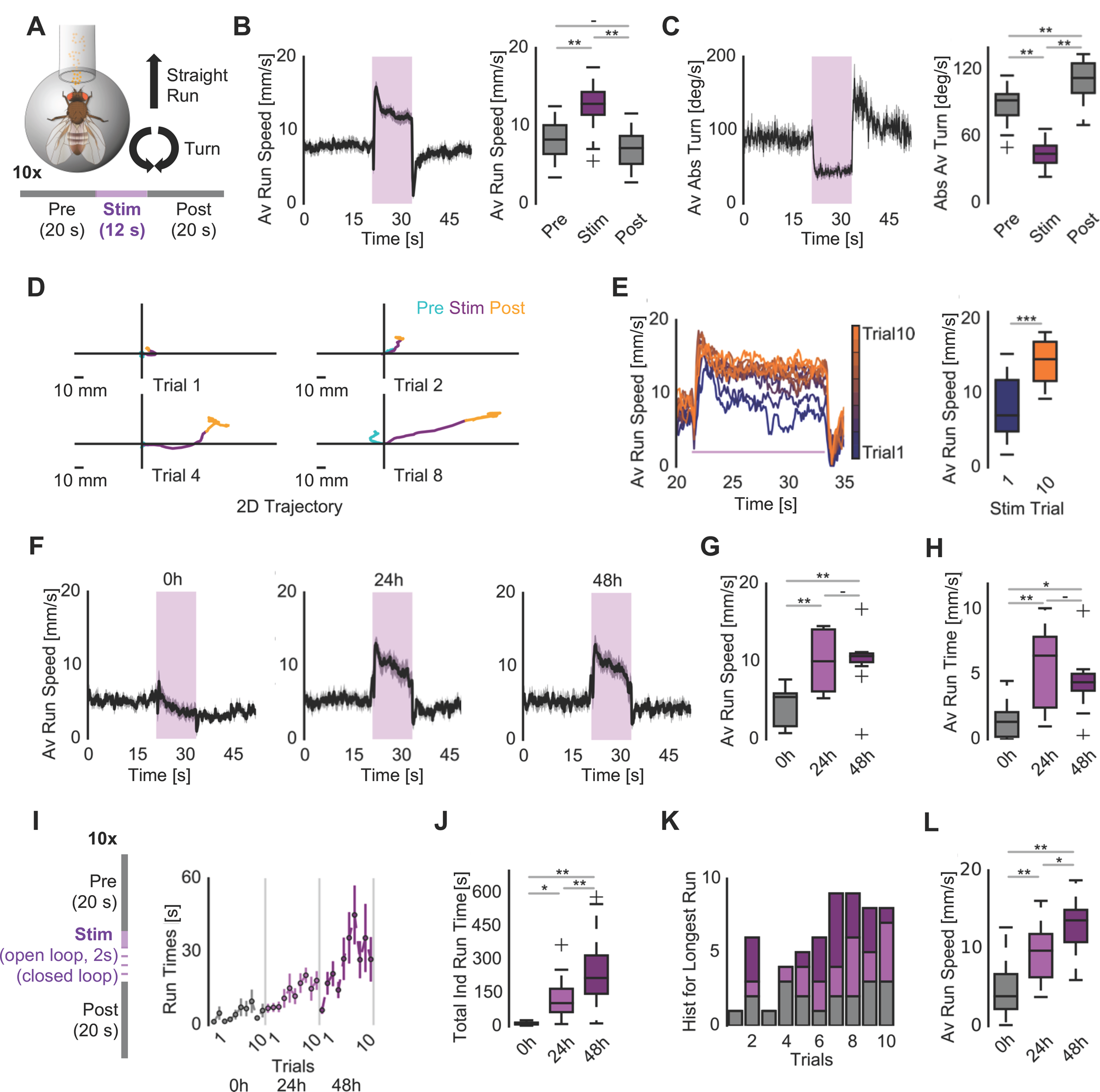
Hungry flies show increasing perseverance to repeated food odor exposure in lieu of a food reward. (A) Spherical-treadmill assay for olfactory stimuli. A tethered starved fly was repeatedly exposed to a 12 s vinegar odor plume for 10 trials during each experiment. Each trial was recorded for 52 s. The inter-stimulus periods were pseudo-randomized in time (60 ± 2-20 s) to reduce odor onset predictability. The locomotor behavior was processed in two cardinal directions, runs and turns, at 10 Hz. Running speed is calculated as the forward locomotion in the odor direction, whereas turning speed is measured as summation of absolute lateral displacement over time. (B) Left: Average running speed (mm/s) of 18 wild-type Canton S strain flies under repeated vinegar exposure for 10 trials. Shaded areas represent the odor exposure duration. On average, flies showed an increased running speed towards the odor source throughout odor presentation. Right: Average running speeds of flies during vinegar exposure were significantly higher compared to the speeds observed during pre- and post-stimulation periods (n=18, one-way ANOVA for correlated samples with Tukey’s HSD post hoc analysis, whiskers represent an extension by 1.5 inter-quartile range). (C) Left: Average absolute turning speed (deg/s) of 18 flies under repeated vinegar exposure for 10 trials. The average turning speed over time and frequency decreased for flies during odor exposure. Right: The absolute turning speed under vinegar was significantly lower than the turning speed recorded in pre- and poststimulation periods. Turns after odor plume loss indicate that flies performed local searches upon loss of odor stimuli, as post-stimulation turning rates were higher than in the pre-stimulation periods (n=18, one-way ANOVA for correlated samples with Tukey’s HSD post hoc analysis). (D) Reconstructed 2D trajectory of a representative fly during single trials. The pre- and post-stimulation behavior for 12 s were depicted in cyan and yellow, respectively, and odor-tracking behavior for 12 s in purple. All trials aligned at the origin for the odor onset. Over trials, the fly showed an increased perseverance in running. The locomotion trajectory data were plotted against pseudo-cardinal coordinates and smoothened with Butterworth filter for visualization. (E) Left: Average running speed of 18 wild flies over time for each of the individual 10 trials. In early trials, flies quickly reverted to the basal pre-stimulation running speeds after initial acceleration at the odor onset. In later trials, flies showed significantly higher perseverance and ran for longer periods of time. Right: Comparison of average running speed between trial 1 and 10. Flies ran on average faster in the last trial compared to the first trial (n=18, paired T-test). (F) Average running speed of fed and hungry flies during repeated vinegar exposure. While fed flies did not track vinegar plumes, flies starved for 24 and 48 hours persistently tracked vinegar. (G) Average running speeds of individual hungry flies during vinegar exposure were significantly higher than fed flies. Starvation level did not alter running speed (n=10/10/11, one-way ANOVA with Tukey’s HSD post hoc analysis). (H) Average running bout lengths in seconds for fed and starved flies. Average running time was calculated from the initial running bouts after odor contact for all trials. When forward running speed was 0 mm/s for 100 ms, the running bout was considered to be terminated. Hungry flies ran longer during vinegar exposure (n=10/10/11, one-way ANOVA with Tukey’s HSD post hoc analysis). (I) Left: Schematics for the closed-loop assay. The closed-loop trials started with a fixed 2 s odor exposure (open-loop) after which the odor channel remained open as long as the animal’s running speed was higher than 0 mm/s (closed-loop). Right: Average running bout times with SEM during closed-loop odor exposure for differentially food-deprived flies over ten trials (n=20/18/19). (J) Average summed running bout times during 10 trials for all groups in closed-loop experiments under vinegar exposure. Longer starvation drove stronger perseverance (n=20/18/19, one-way ANOVA with Tukey’s HSD post hoc analysis. Data were filtered against non-linearity and outliers were removed from the calculation. For outliers, Iglewicz and Hoaglin’s robust test for multiple outliers (Z score ≥ 3.5) was used). (K) Histogram for trials in which flies performed respective longest running bouts throughout 10 trials. The longest run bouts were distributed over trials. For 24 hours starvation, the most frequent peak was observed in the 8^th^ and 10^th^ trials, whereas 48h starvation experimental group reached its peak at the 7^th^ trial. Satiated flies exhibited a random distribution (n=20/18/19). (L) Average running speed of flies during vinegar exposure in the closed-loop paradigm. Longer starvation duration led to increased running speed (n=20/18/19, one-way ANOVA with Tukey’s HSD post hoc analysis, filtered with Iglewicz and Hoaglin's robust test for multiple outliers). For all analyses, statistical notations are as follows: ‘ - ’ > 0.05, ’ * ‘ p < 0.05, ‘ ** ’ p < 0.01, ‘ *** ‘ p < 0.001.

It is possible that any change in the fly’s environment, including a change in airflow or wind, induces forward running. To test this, we analyzed the behavior of mutants of the essential olfactory co-receptor *Orco,* which is required to detect most components of the vinegar odor (Semmelhack and Wang, 2009). *Orco* mutant flies showed a significantly reduced reaction to the vinegar stimulation as compared to their heterozygous controls and hardly accelerated upon stimulus onset (Fig. S1.2A,C). Importantly, these mutants also did not display a similar increase in speed or running time from trial 1 to 10 as observed in control animals (Fig. S1.2B). Loss of Orco, nevertheless, had no significant effect on the animals’ baseline speed before the stimulus (Fig. S1.2D,E). These results suggest that the animal’s reaction depends on the detection of the olfactory stimulus, and indicate that mere practice time on the ball setup do not explain the observed behavior and increase in speed and perseverance.

In addition, we asked whether the valence of the stimulus was decisive for odor tracking behavior in our assay. Previous work looking at turning behavior towards an odor source in a ball setup did not find a difference between the behavior elicited by appetitive as compared to aversive odorants (Gaudry et al., 2013). However, in our assay, frontal stimulation with the highly aversive odorant, CO_2_, led to the opposite behavior compared to the behavior elicited by vinegar (Fig. S1.2F-I). In comparison to a frontal ambient air stimulus, flies slowed down and significantly increased their turning to left and right consistent with odor aversion or an escape response (Fig. S1.2G,H). A similar avoidance behavior was observed in a tethered flying fly assay with frontal odor stimulation (Badel et al., 2016).

Together, these data reveal that food search, or in more general terms goal-seeking behavior, evolves over time, and they indicate that flies work harder in the absence of the expected reward. Furthermore, flies show opposite behaviors for attractive as compared to aversive stimuli. Therefore, this assay provides novel insights into olfactory behavior that go beyond the classical choice assays such as the T-maze or trap assay.

### Tracking perseverance depends on hunger state

Our data show that flies become increasingly persistent over time in odor tracking even in the absence of a reward. What regulates this persistency, which is costly for the animal as it drains energy stores significantly? In all (healthy) animals, the interest in food is regulated by their need to acquire calories and nutrients. This was also evident in this assay: the fed fly did not show any perseverance in food odor tracking in spite of a strong onset and offset behavior (Fig. 1F). By contrast, 24 and 48 h starved flies showed strong tracking behavior and ran continuously on average ~5 s of the 12 s of odor stimulation longer and faster as compared to fed controls (Fig. 1G,H, S1.1 I-L). To better investigate the influence of hunger on the fly’s perseverance, we changed the assay from an open loop to a closed loop configuration, and allowed the fly to control the offset of the odorant by stopping to run (Fig. 1I). The onset of odor stimulation was determined by the experimenter. A stop was defined as 0 mm/s movement for at least 100 ms. As expected, fed flies tracked the odor for 13.3 s, while 24 h starved flies followed the odor for 122.4 s, on average (Fig. 1J). Interestingly, 48 h starved flies showed a much higher perseverance than 24 h starved animals, and tracked the odor for 248.2 s in a single trial, on average (Fig. 1I,J). It is unlikely that the increase in running times is due to improved motor skills, because the increase was not linear or strictly continuous over trials 1-10, but instead among all trials the longest trials of each fly were distributed across all trial numbers including trial number 2 (Fig. 1K). Not only the tracking time but also tracking speed depended on starvation time with 48 h starved animals running on average faster than 24 h starved flies during odor stimulation (Fig. 1L). These results demonstrate that starvation specifically and gradually changes the animal’s perseverance and effort to reach its goal.

### Dopaminergic neuron-provided feedback is necessary for perseverance

A hungry animal following the scent of food expects to find this food eventually. In the present context, the animal remains unrewarded for 10 trials, but instead of giving up the animal increases its efforts. How is the lack of an expected reward translated into perseverance and which neurons represent it? Based on prior evidence discussed in the introduction that DANs are involved in signaling a reward prediction error, we tested whether DANs were involved as reinforcers of behavior. Two major subsets of DANs previously implicated in odor-guided behavior exist in the fly brain: the protocerebral anterior medial (PAM) and the protocerebral posterior lateral (PPL1) cluster (Aso et al., 2014a). In addition, a number of smaller DAN subsets exist in the fly brain including PPL2ab and PPL2c neurons (Mao and Davis, 2009). While the PAM cluster appears to mediate positive experiences during appetitive classical conditioning, PPL1 neurons transmit physically painful stimuli such as electric shock or heat to the fly’s learning center, the MB (Burke et al., 2012; Galili et al., 2014; Liu et al., 2012). The transgenic line *TH-Gal4* labels all cells in the PPL1 and PPL2ab clusters as well as a small subset of the PAM DANs. Thermogenetic activation of these ‘TH+’ neurons can substitute an aversive stimulus and induce negative memories (Aso et al., 2012).

Inactivation of TH+ neurons by overexpression of the temperature-sensitive dominant mutant of dynamin, *shibire* (Kitamoto, 2002), under the control of TH-Gal4 *(TH>shi^ts1^)* changed the fly’s behavior significantly (Fig. 2). The flies showed a reduced average speed during the stimulus and non-stimulus phases over all 10 trials compared to control (Fig. 2A-C). They spend significantly more time turning instead of heading straight on (Fig. 2D,E,I, Fig. S2A,B). Nevertheless, *TH-shi^ts1^* flies did accelerate in response to odorant and the difference between their average speed during pre- and during stimulus phase (i.e. normalized speed) was not different from that of controls (Fig. 2G, Fig. S2D). Similarly, their average speed during the first stimulus phase was also not different from controls (Fig. 2B). By contrast, the increase in straight running speed in response to odor between their first response (first trial) and their last response (last trial) during the stimulus phase was significantly lower (and not different from 0) in *TH-shi^ts1^* animals as compared to controls (Fig. 2F). This strongly suggests that the animals did not augment their behavior as controls did in response to repeated non-rewarded trials. In spite of this, *TH-shi^ts1^* animals spent about the same time being in motion as compared to their genetic controls (Fig. 2H, Fig. S2C). Notably, *Orco* mutants (see Fig. S1.2B) showed a highly similar reduction in average speed and speed increase over trials indicating that olfactory input and the subsequently induced running activity might trigger a DAN-mediated reinforcement signal (Berry et al., 2015; Cohn et al., 2015). It is well known in flies that a permanent lack of dopamine affects motor activity and startle behavior (Riemensperger et al., 2011). Nevertheless, the resemblance of *Orco* mutants and *TH-shi^ts1^* flies and the similar relative acceleration of test and control groups between pre- and stimulus-phase of each trial, support the interpretation that the lack of perseverance and augmentation in tracking behavior rather than primary motor deficits underlie the observed phenotypes. In contrast to TH+ neuron inactivation, inactivation of all PAM neurons with the line *58E02-Gal4* driving *UAS-shi^ts1^* had no effect on the fly’s tracking behavior and perseverance as compared to control flies (Fig. 2A-I). These data support the hypothesis that a subset of DANs, in particular those involved in aversive memory formation, are required as reinforcing signals to drive increased behavioral performance with every non-rewarded trial. Furthermore, they indicate that the negative experience of not finding an expected food reward, i.e. a reward prediction error, is encoded by the same neurons as painful physical experiences.

**Fig. 2:**
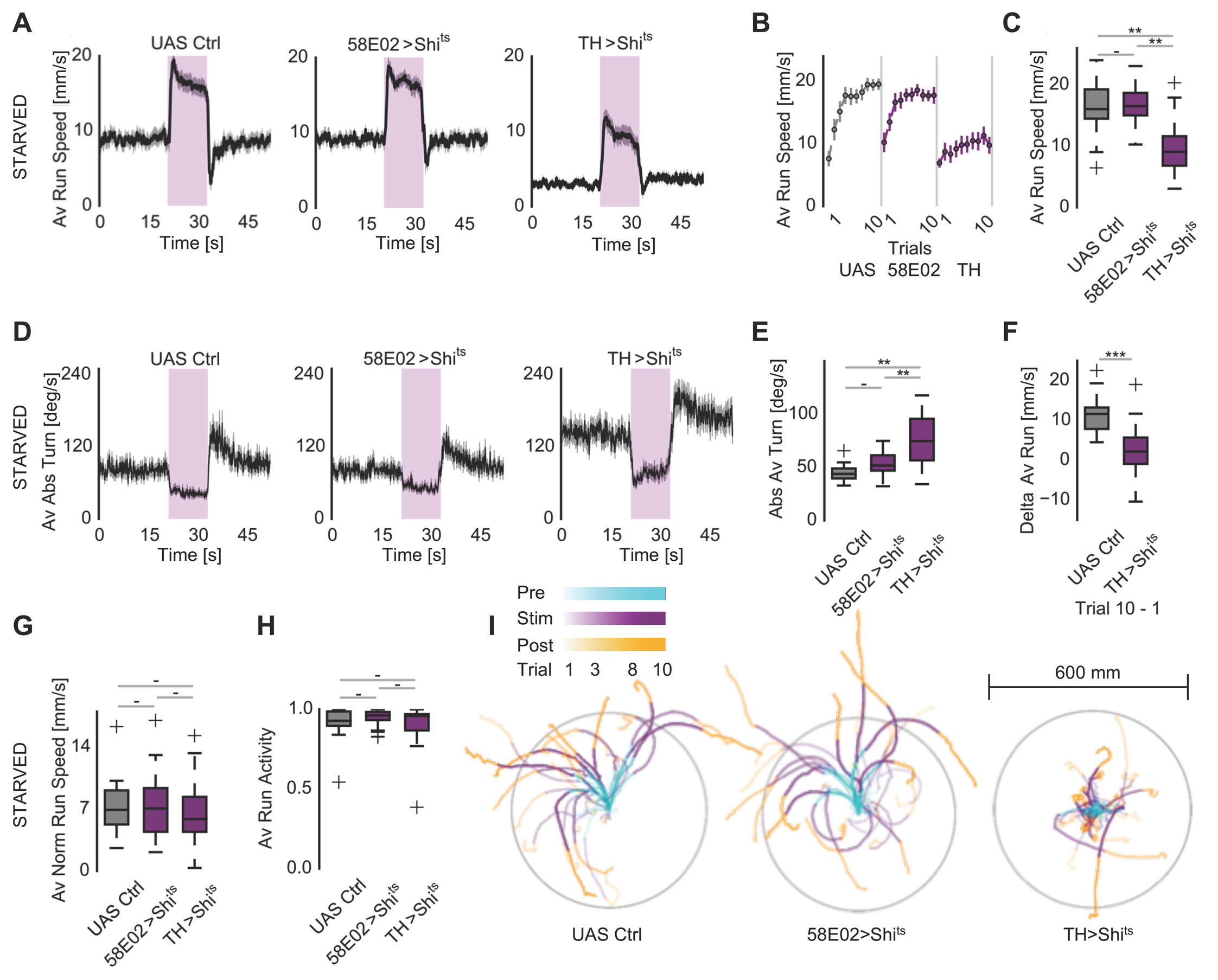
Starvation state-dependent perseverance in odor tracking depends on dopaminergic feedback. (A) Average running speed of hungry flies (24 h starved) with inactivated output of either PAM DANs *(58E02-Gal4>UAS-Shi^ts1^)* or TH+/PPL1 DANs *(TH-Gal4>UAS-Shi^ts1^).* While heterozygous control flies *(+>UAS-Shi^ts1^)* and flies with synaptic output inhibited PAM neurons showed strong and perseverant vinegar tracking behavior, TH+/PPL1 neuron output inhibition strongly reduced average running speed over 10 trials. (B) Average running speed over 10 trials for hungry flies (24 h starved) with inactivated output of PAM DANs *(58E02-Gal4>UAS-Shi^ts1^)* or TH+ DANs (*TH-Gal4>UAS-Shi^ts1^)* compared to control. (C) Quantification of Average running speed of hungry flies (24 h starved) with inactivated output of either PAM DANs *(58E02-Gal4>UAS-Shi^ts1^)* or TH+ DANs *(TH-Gal4>UAS-Shi^ts1^)* compared to control. (D) Average absolute turning behavior of hungry flies (24 h starved) with inactivated output of either PAM DANs *(58E02-Gal4>UAS-Shi^ts1^)* or TH+ DANs *(TH-Gal4>UAS-Shi^ts1^)* compared to control. Flies with inactivated TH+ DANs show a significantly increased turning behavior. (E) Quantification of absolute turning behavior of hungry flies (24 h starved) with inactivated output of either PAM DANs *(58E02-Gal4>UAS-Shi^ts1^)* or TH+ DANs *(TH-Gal4>UAS-Shi^ts1^)* compared to control. (F) Difference between the speeds at trial 10 versus trial 1. While controls run clearly much faster on average during odor stimulation at trial 10, TH+/PPL1 DAN inactivation strongly diminishes this effect and these flies do not show a significant acceleration over trials. (G) Average normalized running speed upon transiently blocking DAN output. In the absence of PAM cluster *(58E02-Gal4>UAS-Shi^ts1^)* or TH+ positive *(TH-Gal4>UAS-Shi^ts1^)* dopaminergic neuron output, average running speeds during vinegar exposure were comparable to the genetic control after baseline normalization to the pre-stimulation phase suggesting that all groups accelerate equally in the presence of the odor stimulus. (H) All groups also show a similar level of overall activity (time in motion (>0 mm/s)) during stimulus phase. (I) 2D representation of tracks of 7 randomly chosen flies/condition for *UAS-Shi^ts1^* controls, *58E02>Shi^ts1^*, and *TH>Shi^ts1^* experimental groups. The lighter the shading of the trial, the earlier in the experiment. Trials 1,3,8, and 10 were plotted for all 7 flies. To facilitate graph interpretation a grey circle was drawn at 300 mm from the starting point of the fly. The absolute heading direction displayed is irrelevant as the odor was always kept in front of the animal. Note that inactivation of the synaptic output of TH+ neurons strongly reduces heading straight and increases turning. For all analyses, statistical notations are as follows: ‘ - ’ > 0.05, ’ * ‘ p < 0.05, ‘ ** ’ p < 0.01, ‘ *** ‘ p < 0.001.

### Specific MB output neurons are involved in food odor tracking

DANs form synapses with Kenyon cells (DAN>KC) in a highly MB lobe region-specific manner (Aso et al., 2014a). Recent data have shown that DANs connect directly to the so-called MB output neurons (MBONs; DAN>MBON), and are also innervated by KCs themselves (KC>DAN) (Eichler et al., 2017; Takemura et al., 2017). In addition, some MBONs reconnect and innervate KCs (MBON>KC). The degree of the individual MBON’s output is subject to neuromodulation by dopaminergic, serotonergic, GABAergic and other neuron types. We next addressed, whether MBONs are required for food odor tracking and if it were the case, which MBONs might be modulated - directly or indirectly - by the feedback of DANs. Using screening of Split-Gal4 lines that targeted MBONs for a phenotype in attraction to vinegar with the T-maze assay, we identified the line MB112C, which expresses in the so-called MVP2 or MBON-Y1pedc>a/p neurons (Aso et al., 2014b) (Fig. S3.1A-D). Inactivation of MVP2’s synaptic output using overexpression of the transgene *shi^ts1^* significantly reduced olfactory tracking behavior (Fig. 3A-C, Fig. S3.2A). In order to test whether activation of MVP2 was sufficient to induce odor tracking in the fed fly, we expressed the red-shifted channel rhodopsin CsChrimson under the control of MB112C *(MB112C>CsChrimson),* and stimulated the flies with the usual 12 s odor stimulus, but overlapped it 2 s after odor onset with a 10 s pulsed red-light stimulus (Fig. 3D-F). Activation of MVP2 using this optogenetic protocol in fed flies indeed induced significant tracking in the presence of an odor stimulus (Fig. 3D-F, S3.2B,C). The same manipulation did not increase tracking in starved flies indicating that MVP2 is already active in the food-deprived animal during odor tracking (Fig. S3.2D,E) (Perisse et al., 2016). Furthermore, light alone in the absence of frontal odor stimulation did not induce more forward running compared to CsChrimson controls or wildtype flies (Fig. S3.2F-H). These data show that MVP2 is necessary and sufficient for food odor tracking in the present context.

**Fig. 3:**
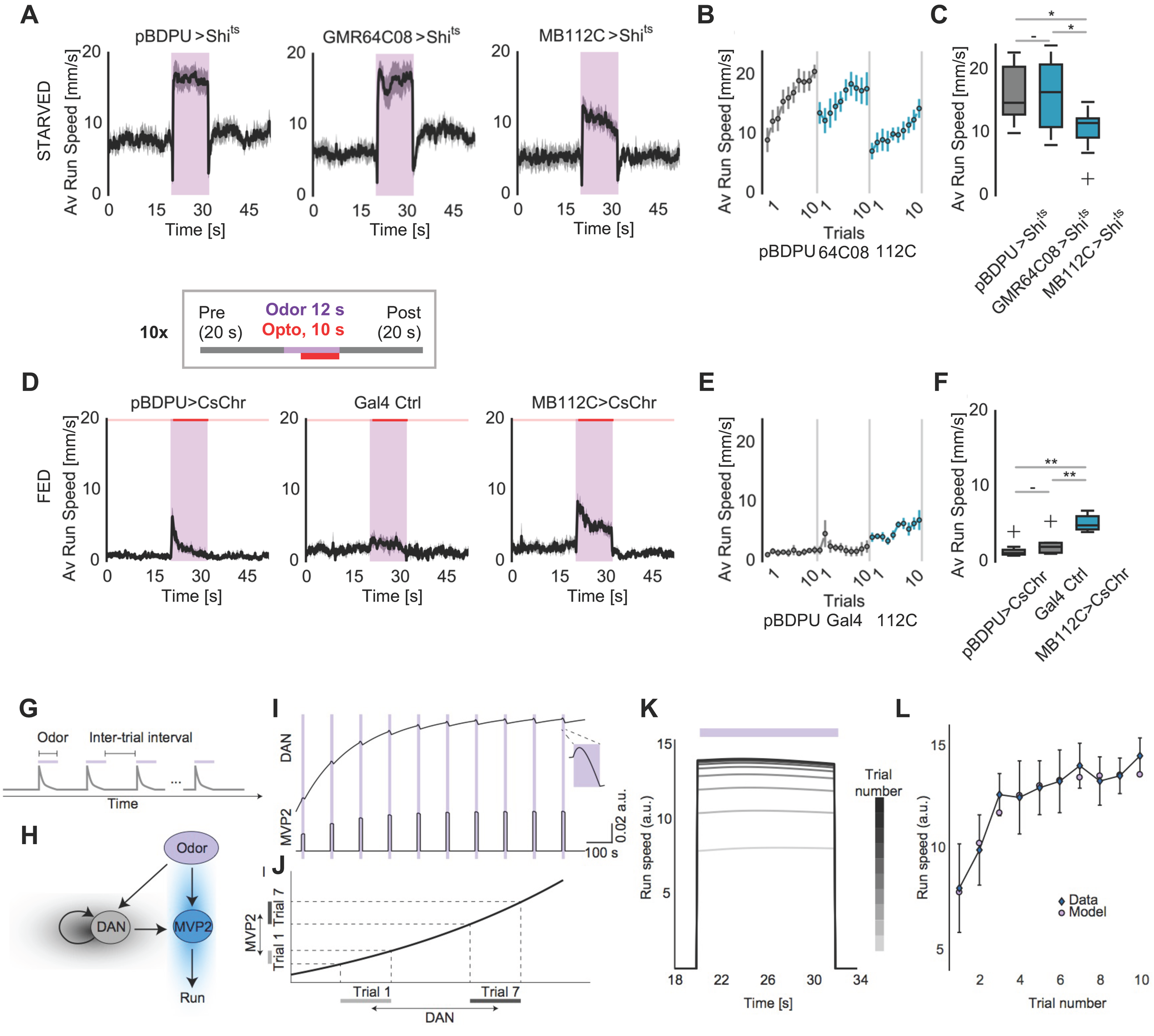
A specific mushroom-body output neuron induces odor tracking. (A-C) Average running speeds upon blocking synaptic output of MVP2/MBON-γ1pedc>αβ and mushroom body γ Kenyon cells at non-permissive temperature through overexpression of dynamin mutant UAS-Shibire^ts1^. GMR64C08-Gal4 labels all mushroom body γ Kenyon cells. The heterozygous control was empty-Gal4 *(pBDP-Gal4U¾UAS-Shibire^ts1^).* While blocking all γ Kenyon cells did not alter vinegar tracking, the synaptic input from the MVP2 neurons was necessary for odor tracking behavior (n=10/10/10). (D-F) Average running speeds during acute MVP2 activation with CsChrimson *(MB112-Gal4¾UAS-CsChrimson)* in fed flies, comparison to controls *(pBDP-Gal4U>UAS-CsChrimson* and Gal4 Ctrl: *MB112-Gal4¾+).* Concurrent odor application and MVP2 activation with light induced odor tracking in fed flies (n=7/6/7). To reduce a light onset startle effect, a constant low intensity background of 617nm light (0,15 pW/mm^2^) was used throughout the entire experiment. (G-K) Model captures the persistence of running speed in the activity of DANs. (G) Repeated odor presentation separated by odor-free intervals in the model. During each trial, odor presentation is represented as a step increase in the stimulus for 12 s (purple bars); during inter-stimulus intervals the stimulus is zero for 105 s. The input to the DANs is odor-dependent and in the form of a decaying stimulus to mimic, for instance, PN-transformed odor response (Fig. S3.3E, gray). Bottom: Circuit diagram. In the presence of odor, MVP2 drives running behavior (blue shading). During inter-stimulus intervals, DAN activity accumulates negative experience due to lack of reward (gray shading). This activity modulates MVP2 during the next odor presentation. (H) Simulated DAN activity for 10 consecutive trials separated by inter-stimulus intervals. During odor presentation (purple bar), DANs integrate the transient PN responses. Between stimuli, DANs accumulate the reward prediction error from the previous odor presentation with a time constant that is slower than the inter-stimulus interval. This generates DAN activity that persistently increases with trial number. MVP2 integrates the odor with a faster time constant and is modulated by the DANs, but only during odor presentation. (I) A nonlinearity transforms MVP2 output into speed, demonstrating that gradually increasing DAN activity more strongly activates MVP2 in later trials. (J) Running speeds per trial generated by the model (compare to Fig. 1E). The model captures the persistent behavior. (K) Comparison between the average speed per trial obtained by the fitted model and the data (mean +/- standard deviation of Fig. 1E). Model parameters (see Table 1 for sensitivity analysis): *α = 171,b* = 8*,θ* = 0.25*,α_y_* = 1,5 = *41, k* = 0.001, A = 0.002, τ = 3, *R = 0,r =* 0.

MVP2 is an inhibitory neuron with dendrites in the MB γ1 peduncular region and with axons innervating the α and β lobes (Aso et al., 2014a). Given our data above suggesting that DANs provide error feedback to the animal, which conceivably translates into increased perseverance to find the expected food, we asked whether KCs are involved in the observed behavior. We, therefore, tested whether γ-type Kenyon cells were required for odor tracking by inactivating their synaptic output again with shi^ts1^*(GMR64C08>shi^ts1^)*(Fig. 3A-C, 3.2A). This manipulation, however, had no effect showing that γ-type Kenyon cell output is not important for food odor tracking in contrast to the γ-MBON MVP2. Moreover, inactivation of all KC synaptic output using a broader line *(MB10B>shi^ts1^)* also did not affect odor tracking of the hungry fly on the ball (Fig. S3.3A-D). While these data do not fully rule out a function of KCs that was not uncovered under the current conditions, they support a role for a direct interaction between DANs and MBONs.

### Modeling the role of dopamine in perseverance

To further investigate the possible neural mechanisms underlying the observed perseverant behavior in the continued presence of odor but lack of reward, we proposed a minimal circuit model of DANs and MVP2 (Fig. 3G). We focused on these two populations for three reasons. First, we already showed that broad inactivation of DANs (TH+ neurons) in starved flies decreases perseverance (Fig. 2), which indicates that these neurons have a determinant role. Second, inactivation of MVP2 reduced the hungry fly’s olfactory tracking (Fig. 3A-C), suggesting that information from the DANs modulates MVP2 activity or output. In fact, since MVP2 has dendrites in the MB γ1 peduncular region and axons in the α and β lobes, it could be modulated by several DAN types in the MB peduncle (input level; e.g. MP1) and α and β lobes (output level; e.g. V1 and V2) (Aso et al., 2014a; Perisse et al., 2016; Takemura et al., 2017). Third, recent connectomics data have demonstrated a direct connection from DANs onto MBONs arguing that DANs modulate behavior without KC involvement (Eichler et al., 2017; Takemura et al., 2017). To mimic the experimental setup, we simulated 10 trials of 12 s of external input (e.g., odor), separated by inter-stimulus intervals of 105 s (Fig. 3G; see Methods). We modeled MVP2 activity only during odor presentation, because exogenous MVP2 activation does not induce running behavior in the absence of odor (Fig. S3.2C). Since *Orco* mutant flies showed lack of perseverance (Fig. S1.2B), we assumed that the input to DANs is odor-dependent. We used a slowly decaying odor-dependent stimulus to drive MVP2 during odor presentation (Fig. 3G, blue shading) based on the slow dynamics observed during *in vivo* calcium imaging of olfactory projection neurons (PNs) that connect ORNs to higher brain centers (Figure S3.3E). This stimulus generated a transient ‘bump’ in the activity of the DANs, which was followed by a gradual decrease (Fig. 3H). During inter-stimulus periods we hypothesized that the activity of DANs will continuously increase due to the accumulation of negative experience from the lack of the predicted reward (Fig. 3G, gray shading). Therefore, the model DANs accumulate a reward prediction error signal generated during repeated odor presentation without reward, which then modulates the activity of MVP2 at the input or output level and directs the animal to run faster on each trial (Fig. 3H).

We postulated that the increase in DAN activity between trials is determined by a long time constant, which captures the speed of accumulation of non-rewarded experience: longer time constants induce a more perseverant search for food. To determine this time constant, we fit our model output (upon transforming MVP2 activity to speed) to the measured running speeds (Fig. 1E; Methods). We consistently obtained values longer than the inter-stimulus interval (162.6±13.0 s vs. 105 s; Table 1 in Methods), providing strong evidence that DAN activity is outlasting the inter-stimulus period, and therefore can indeed impact and reinforce behavior at the next trial. If this time constant were shorter than the inter-stimulus interval, it would mean that the ‘memory’ of the latest negative experience would not be available in the next trial, and thus persistence would not be observed. Such long-time constants could be the result of feedback loops, which are common in the mushroom body, where DANs and MBONs innervating the same compartment have the potential to form recurrent connections (Aso et al., 2014a; Ichinose et al., 2015; Zhao et al., 2018).

To transform MVP2 output into running speeds, activity was passed through a monotonically increasing nonlinearity, which scales neural output to capture the measured speeds (Fig. 3J; Methods). Therefore, the gradual increase in DAN activity across trials shifts MVP2 activity (and hence running speed) to higher values, whereby MVP2 becomes more activated in later trials, driving faster running (Fig. 3H). The persistent behavior captured by the model with speeds that increase as a function of trial number indeed corresponds well to the persistent behavior measured in the flies (Fig. 3K).

Our model shows that the accumulation of reward expectation can generate persistent behavior, and that the accumulated prediction error signal mediated by the DAN population is a key element. It also predicts that the time constant for DANs is longer than the inter-stimulus interval, consistent with the idea of persistent DAN activity arising from recurrent feedback loops in the mushroom body (Ichinose et al., 2015; Zhao et al., 2018).

### Sequential antagonistic behaviors distinguish goal seeking from reaching the goal

In most healthy animals, hunger motivates food search. If the expected food is not found, the animal will search longer at the next occasion to find it. This perseverance depends on dopaminergic signaling as shown above. What happens, however, if the food search is successful? Once food has been found, search behavior should be suppressed for the animal to engage into tasting the food and, if to its liking, ingesting it. Hence, a successful food search can be divided into at least three antagonistic steps: (i) search and long-distance tracking or hunting (e.g. olfactory cue), (ii) reaching the food and short-distance evaluation (e.g. gustatory cue), and (iii) ingestion and post-ingestion reward (e.g. internal sensory cues, internal sugar levels). In our assay, we observed tracking in response to an olfactory cue. Reaching the food source and ingesting the food should, by contrast, suppress locomotion (Mann et al., 2013; Thoma et al., 2016). Presumably, this should also happen despite the continued presence of the odorant, in order to allow the fly to feed. To test this prediction, we expressed CsChrimson and optogenetically activated two different sets of sugar neurons as a proxy for food and food ingestion 2 s after the onset of the vinegar stimulus (Fig. 4A). First, activation of exclusively peripheral (labellar and tarsal) gustatory receptor (Gr) 5a-expressing sweet taste neurons was employed to mimic the second step in a food search; reaching the food and evaluating its taste (Thoma et al., 2016). Next, Gr43a is not only expressed in peripheral taste neurons, but has also been shown to function as internal sugar sensor in the brain (Miyamoto et al., 2012). Gr43a is also among the taste receptors expressed in enteroendocrine cells in the fly gut (Park and Kwon, 2011). They were therefore activated to mimic a post-ingestion reward. Gr5a activation led to an immediate stop of odor tracking suggesting that animals that reach a food source cease searching for food (Fig. 4 A-C, Fig. S4A-C). Interestingly, although stimulation with light continued for 10 s, flies quickly resumed odor tracking and reached their full speed again at the end of each trial (Fig. 4A,C). By contrast, Gr43a activation lastingly reduced the fly’s tracking behavior consistent with sensing the rise of internal sugar levels upon successful ingestion of a sweet food (Fig. 4A-C). Different from flies with activated Gr5a neurons, flies with active Gr43a neurons did not accelerate again after a few seconds of light stimulation, but instead remained at the same slow speed till the end of the stimulation period (Fig. 4C). In this regard, it is interesting to note that Inagaki et al. have previously reported that prolonged optogenetic activation of sugar taste neurons (i.e. Gr5a) only transiently induces proboscis extension, which ceases much before the end of the light stimulation (Inagaki et al., 2014).

**Fig. 4:**
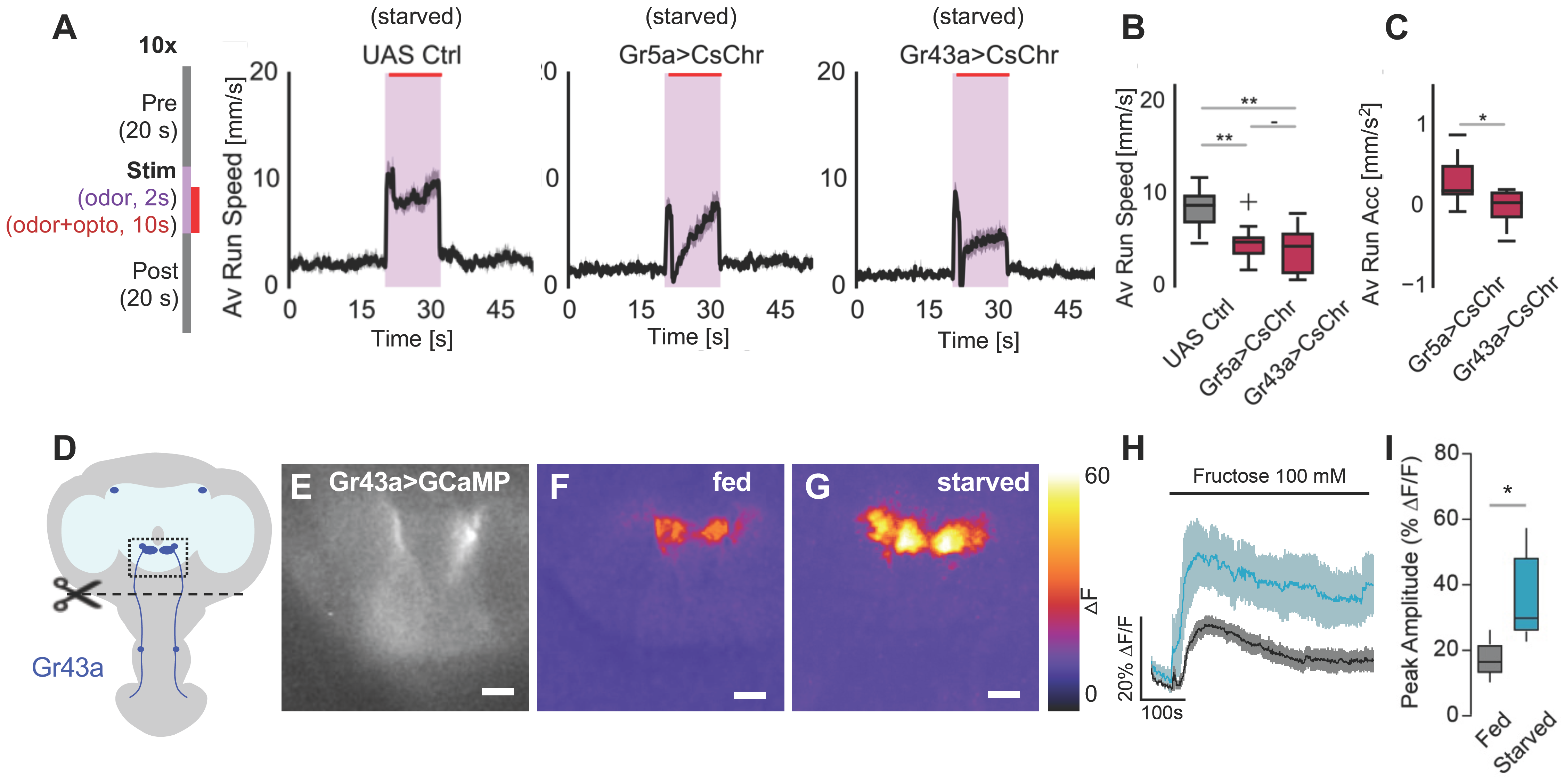
Internal and peripheral sugar taste neuron activation interrupts odor tracking. (A) Left: Schematics of the concurrent odor and optogenetic-activation protocol. A 617 nm high-power single LED was used for optogenetic neuronal activation after 2 s of odor delivery in each trial. Right: Activation of peripheral and internal sugar taste neurons to mimic a food reward. *Gr5a>UAS-CsChrimson* flies express CsChrimson in peripheral sweet taste receptor neurons. *Gr43a>UAS-CsChrimson* flies drive CsChrimson expression in peripheral and pharyngeal sweet taste and internal sugar receptor neurons. UAS-Ctrl is the genetic background control with an empty Gal4 transgene *(pBDP-Gal4U>UAS-CsChrimson).* (B) Average running speed during simultaneous odor and goal presentation. Optogenetic activation of Gr5a and Gr43a positive neurons significantly reduced odor tracking speed. Statistical analyses were performed for the period of simultaneous odor and optogenetic stimulation (n=10/10/10, one-way ANOVA with Tukey’s HSD post hoc analysis). Note that Gr5a activation does not lastingly suppress odor tracking. (C) Average acceleration during simultaneous odor and mimicked taste or nutrient presentation. Gr5a activated flies accelerated quickly after the optogenetic onset, while Gr43a flies remained slow or stopped (n=10/10/10, one-way ANOVA with Tukey’s HSD post hoc analysis). (D-I) Ascending axons from pharyngeal Gr43a neurons respond to fructose *ex vivo.* (D) Fly head schematic showing the location of pharyngeal Gr43a neurons and their projections to the SEZ in the central brain. (E) Grayscale image showing the basal expression of *Gr43a>GCaMP6f* in the SEZ in an *ex vivo* preparation. (F-G) representative pseudo-colored images showing the GCaMP6f intensity change in response to bath application of fructose on explant brains from fed and 24h starved flies (100 mM, scale bars 20 μm). (H) Time course of GCaMP6f intensity changes in response to fructose (mean ± SEM; n=8/5). (I) Fluorescence peak intensity change in explant brains from fed and 24h starved flies (unpaired T-test). For all analyses, statistical notations are as follows: ‘ - ’ > 0.05, ’ * ‘ p < 0.05, ‘ ** ’ p < 0.01, ‘ *** ‘ p < 0.001.

We conclude that starvation state is the main motivator for persistent food odor tracking and reaching the attempted goal readily suppresses this behavior transiently or lastingly depending on successful food ingestion and post-ingestion signals.

Since Gr43a neuron activation is sufficient to stop the fly from search for food, we wondered which phase of feeding these neurons detect. In addition to their role in internal sugar sensing, prior work has shown that pharyngeal Gr43a taste neurons connect to interneurons that modulate the actual intake of food and ingestion process depending on internal state and food quality (Yapici et al., 2016). We therefore asked whether peripheral Gr43a neurons would also be sensitive to internal sugar level changes, not limited to taste and feeding itself. To this end, we carried out calcium imaging of the Gr43a-positive axons innervating the SEZ in an *ex vivo* brain only preparation, where these axons would no longer be connected to their pharyngeal cell bodies and dendrites. Indeed, these severed GR43a axons reacted significantly to the bath application of 100 mM fructose (Fig. 4D-I). The response of these axons followed a similar slow dynamic as previously observed for the central Gr43a neurons (Miyamoto et al., 2012), and was significantly slower than the response of peripheral stimulation of Gr43a neurons in the pharynx (LeDue et al., 2015). Furthermore, we observed that this response was modulated by hunger, as Gr43a axons responded significantly better to fructose stimulation in brains from hungry animals as compared to brains from fed animals (Fig. 4H,I). This suggests that Gr43a axons that project from the periphery into the SEZ are sensitive to increasing sugar levels in the lymph and are therefore conceivably involved in all stages (taste, ingestion, and post-ingestion cue detection) of the feeding process.

### Octopamine suppresses tracking in hungry flies

Food tracking, hunting or searching, food evaluation, and food ingestion represent sequential and antagonistic behaviors. Dopamine provides the information that the expected goal has been missed. But how is a positive experience conveyed? And how is sensory information coordinated and prioritized such that a gustatory external or internal nutrient stimulus overrides or enforces odor-stimulated behavior? Successful appetitive olfactory learning (i.e. pairing and odor with a sugar reward) is driven by the other set of DANs, the PAM cluster neurons. Inactivation of PAM DANs, however, did not appear critical in the present assay (see Fig. 2).

Based on previous work, which has elegantly shown that starvation modulates peripheral chemosensory neurons (i.e., olfactory neurons become more sensitive to food odor and thereby facilitate the fly’s food search) (Root et al., 2011), it is formally possible that post-ingestion signals desensitize olfactory neurons and fed flies become, so-to-speak, odor blind. However, with the same stimulation protocol as used in the behavioral assay, we carried out an *in vivo* calcium imaging experiment of olfactory projection neurons (PNs) that connect ORNs to higher brain centers. We found that PNs not only did not adapt, but continued to respond during repeated long-term stimulation; they also responded similarly in starved and fed animals (Fig. S3.3E,F). These results suggest that olfactory stimuli continue to be detected over time and also by fed animals that do not show persistent odor tracking (see Fig. 1F).

Next, we investigated the involvement of OA in the interplay of starvation, food search and food consumption for three reasons. First, as described above, OA was previously implicated in motivation in *Drosophila.* Second, due to their location and projection patterns, a subset of OA neurons, similar to the NE neurons in the NST, appeared suited for communication between nutrient information and higher (olfactory) brain centers. Third, Burke et al. have previously implicated a group of ventral OANs that project towards the MB as signals of taste valence (Burke et al., 2012). We used a transgenic fly line that expresses Gal4 under the control of the Tdc2 promotor *(Tdc2-Gal4)* to express the temperature sensitive channel TrpA1 in OA neurons (Hamada et al., 2008). First, we activated Tdc2+ neurons in a starved animal during the entire experiment by shifting the flies to 30^°^C briefly before and during the experiment. These flies, but not their respective genetic controls showed complete suppression of odor tracking and instead stopped immediately at odor onset instantaneously at the first trial (Fig. S5.1A-E). In addition, the flies accelerated quickly after odor offset and regained speeds similar to controls (Fig. S5.1A). This behavior was also observed in fed flies indicating that OA neuron activation overrides feeding state (Fig. S5.1F,G). By contrast, chronic activation of OA neurons using TrpA1 had no significant effect on the flies’ running speed, when they were stimulated exclusively with ambient air and not food odor (Fig. S5.1H,I).

OANs were previously implicated in appetitive learning as a part of the reward system in *Drosophila* and other insects (Burke et al., 2012; Schroter et al., 2007; Schwaerzel et al., 2003). Based on previous data and our observations, it is possible that OANs mimic either an acute reward (e.g. food taste or post-ingestion signal similar to Gr43a neurons) or a more chronic internal state such as arousal, hunger or satiation (Perry and Barron, 2013). To distinguish between these possibilities, we tested whether also acute activation of OA neurons resulted in tracking suppression by using optogenetic activation of Tdc2 neurons paired with the odor stimulus (Fig. 5A-C, Fig. S5.1J-K). This acute activation *(Tdc2>CsChrimson)* 2 s after odor onset fully suppressed odor tracking and led to immediate and persistent slowing down or stopping, while flies with the same genetic make-up stimulated only with odor and not light behaved exactly like controls (Fig. 5A-C, Fig. S5.1L,M). Importantly, flies also did not increase and even decreased their turning frequency during OA neuron plus odor stimulation (Fig. S5.1N,O). This result strongly indicated that OA neuron activation did not induce an aversive response, as this would have increased turning behavior (see Fig. S1.2). Therefore, OA neurons in the present context might acutely transmit the presence of food and/or a post-ingestion effect rather than a more chronic internal state of arousal or hunger.

**Fig. 5.**
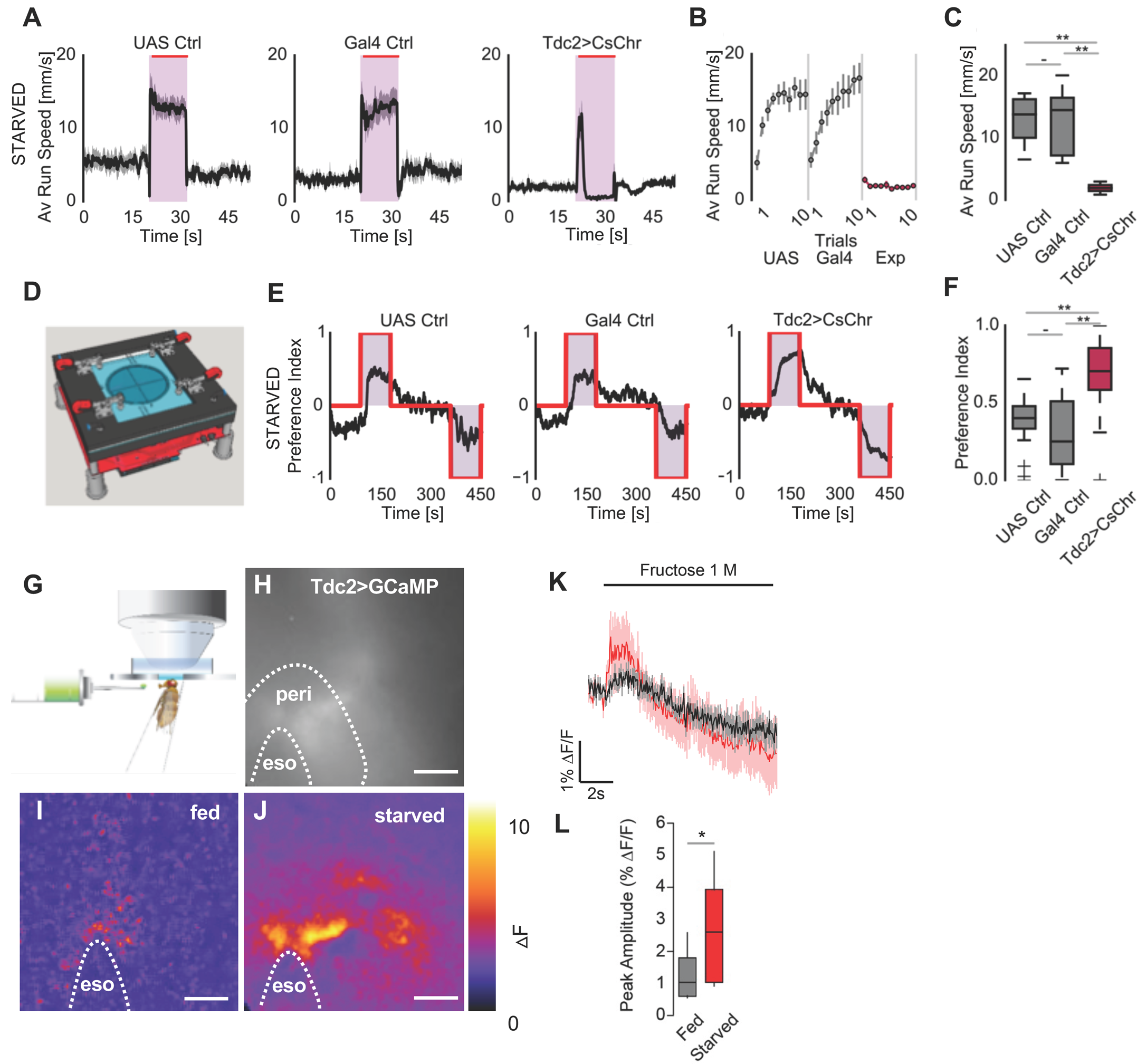
Activation of octopaminergic neurons induces quitting of olfactory tracking. (A) Acute optogenetic activation of octopaminergic neurons. CsChrimson was expressed in octopaminergic neurons by *Tdc2>UAS-CsChrimson* (Controls: UAS Ctrl: *+>UAS-CsChrimson,* Gal4 Ctrl: *Tdc2-Gal4>+).* Acute activation of octopaminergic neurons for 2 s after odor onset instantaneously reduces odor tracking, suggesting that Tdc2+ neurons are required acutely inconsistent with a role in chronic starvation or arousal. (B) Evolution of average running speeds for *Tdc2>UAS-CsChrimson* flies during odor exposure over trials. (C) Average running speed for *Tdc2>UAS-CsChrimson* flies during odor exposure (n=10/10/10, one-way ANOVA with Tukey’s HSD post hoc analysis). (D) Scheme of optogenetic and olfactory behavioral test arena. ~15 flies were loaded into a 10 cm circular arena, where each quadrant could be controlled for combined odor and optogenetic stimulation. A concurrent stimulation paradigm with odor and light was used. After a prestimulation period, a pair of opposing quadrants were switched on for a vinegar odor stream and LED illumination for 90s (red quadrants). Following the inter-stimulus period, the reciprocal pair of quadrants were activated. Fly behavior was analyzed as a preference index, based on the distribution of flies in the last 5 s of each stimulation period ((# flies in stim quadrant - # flies in non-stim quadrant) / (# flies in stim quadrant + # flies in non-stim quadrant)). Due to the calculation method, positive PIs in the first and negative PIs in the second stimulus phase display attraction to the odor quadrant. (E) Preference index over time for vinegar in the arena for control and flies with activated octopaminergic neurons (n=16/16/16). (F) Average preference index during optogenetic activation of octopaminergic neurons under vinegar exposure. Activation of octopaminergic neurons led to significantly more accumulation of flies in the odor quadrants when compared to genetic controls (n=16/16/16, one-way ANOVA with Tukey’s HSD post hoc analysis). (H-M) Octopaminergic neurons respond to proboscis fructose application *in vivo.* (G) Schematic representation of the *in vivo* imaging setup. (H) Grayscale image showing the expression of Tdc2>GCaMP6f in the ventral fly brain (*eso,* esophagus; *peri,* periesophageal zone). (I,J) Representative pseudocolored images showing the GCaMP6f intensity change in response to the application of a drop of fructose solution on the proboscis of fed and 24h starved flies (1 M, scale bars 20 μm). (K) Time course of GCaMP6f intensity changes in response to fructose (mean ± SEM; n=10/9). (L) Fluorescence peak intensity change in fed and 24h starved flies (unpaired T-test). For all analyses, statistical notations are as follows: ‘ - ’ > 0.05, ’ * ‘ p < 0.05, ‘ ** ’ p < 0.01, ‘ *** ‘ p < 0.001.

To gain more evidence that Tdc2+ neuron activation indeed represented something rewarding such as finding food for the animals, we used a custom-built 4-arm olfactory choice assay (Fig. 5D). Optogenetic activation of Tdc2+ neurons in the presence of vinegar odor in the same quadrant resulted in a significantly higher dwell time of the flies in the illuminated quadrants (Fig. 5E,F) suggesting that octopaminergic neuron activation is indeed rewarding.

Interestingly, blocking synaptic output of OA neurons *(Tdc2-Gal4>shi^ts^)* did not show the opposite behavior as compared to OA neuron activation. Instead, flies of this genotype moved less compared to controls; this was true for forward running as well as for turning in either direction (Fig. S5.2E-H). A similar phenotype was observed in flies mutant for TβH, the enzyme required to generate octopamine (Fig. S5.2A-D). While these data are consistent with the previously reported systemic role of octopamine in arousal and motor activity, they indicate that different types of OA/Tdc2+ neurons might play different and potentially contrary roles.

In summary, these results are consistent with a role of octopamine as a mediator of rewarding information such as the taste and the nutritive value of a food. Hence, they are good candidates to promote the behavioral switch between seeking and stopping when having reached a goal.

### Ventral cluster OA neurons are sugar sensitive

Activation of Gr43a and Tdc2+ OA neurons resulted in a similar behavior of the fly. Since Gr43a axons in the SEZ have the potential to respond to internal sugar level changes, we next tested whether ventral cluster OA neurons were sensitive to sugar (Fig. 5G-L). Previous work has shown that likely via direct synaptic connection Gr32a taste neurons responding to male pheromone taste activate Tdc2+ neurons in the SEZ (Andrews et al., 2014). To test, if sugar taste would activate ventral OA neurons, we used *in vivo* calcium imaging and stimulated the flies with a fructose-solution filled feeding pipet held to their labellum (Fig. 5G). Importantly, the proboscis was fixed preventing movement and allowing forced touching of the labellum independent of voluntary proboscis extension. We focused on the region of the brain, where the ventral OANs extend the majority of their dendrites (SEZ and periesophageal zone; Fig. 5K). We observed that Tdc2+ neurons responded to sugar (Fig. 5H-K); and this response was again significantly higher in starved animals as compared to fed animals (Fig. 5L). Similar to our results, previous imaging experiments of Tdc2+ neurons in response to taste (Gr32a-mediated) revealed very small changes in calcium transients (Andrews et al., 2014). In addition, work in honeybee found that a ventral OAN, the so-called VUMmx1 (ventral unpaired medial), which is involved in appetitive memory formation, responds to sugar with a similar timing as compared to our observation (Hammer, 1993). Essentially, VUMmx1 showed initial high-frequency bursting for 1-2 s followed by a longer period of around 30 s of regular lower frequency spikes in response to sugar taste (Hammer, 1993). Such prolonged spiking does not necessarily increase cellular calcium transients. Therefore, and due to photo-bleaching of the indicator, it is possible that a longer lasting low signal was not detected.

Nevertheless, these results strengthened our hypothesis that OANs counteract the animal’s perseverance to keep running after an odorant by conveying sugar reward to the higher brain.

### A specific subtype of OA neurons antagonizes food seeking

The ventral cluster of Tdc2+ neurons contains several types of OANs such as the VUM and VPM (ventral paired medial) type neurons, which again contain different anatomically distinct types of neurons (Busch and Tanimoto, 2010).

To pinpoint the exact neuron(s) capable of suppressing odor tracking in response to food reward, we screened several candidate OA neuron lines that fit our above-mentioned criteria by optogenetic activation (starting 2 s after odor onset) for the phenotype observed upon activation of sugar sensitive GR neurons. Two Split-Gal4 transgenic lines (Aso et al., 2014a), MB113C and MB22B labeling two neuron types of the VPM cluster, VPM3 and VPM4, showed a phenotype that closely resembled activation of Gr43a neurons (Fig. 6A, Fig. 6.1A-C). This included the strong and sustained decrease in odor tracking behavior during light plus odor stimulation (Fig. 6B,C, Fig. S6.1C). By contrast, genetic controls and flies carrying the same transgenes, but which were stimulated only with odor and not light, behaved exactly as wildtype controls flies (Fig. S6.1D-G). More chronic activation of these neurons using TrpA1 thermogenetics during the entire experiment and ca. 60 min before also led to comparable phenotypes without affecting running speed during pre-stimulus phases providing further evidence for a more specific role of these neurons in food seeking behavior (Fig. 6.2A-L). Inhibition of the synaptic output of these VPM neurons did not result in a change in food odor tracking (Fig. S6.3A-D) showing that these neurons are likely not directly involved in the execution of tracking behavior or olfactory processing. In addition, they are either redundant or not part of the OA neurons that are responsible for reduced odor tracking of *TβH* mutants or upon blocking of all Tdc2+ neurons’ synaptic output (see Fig. S4).

**Fig 6.**
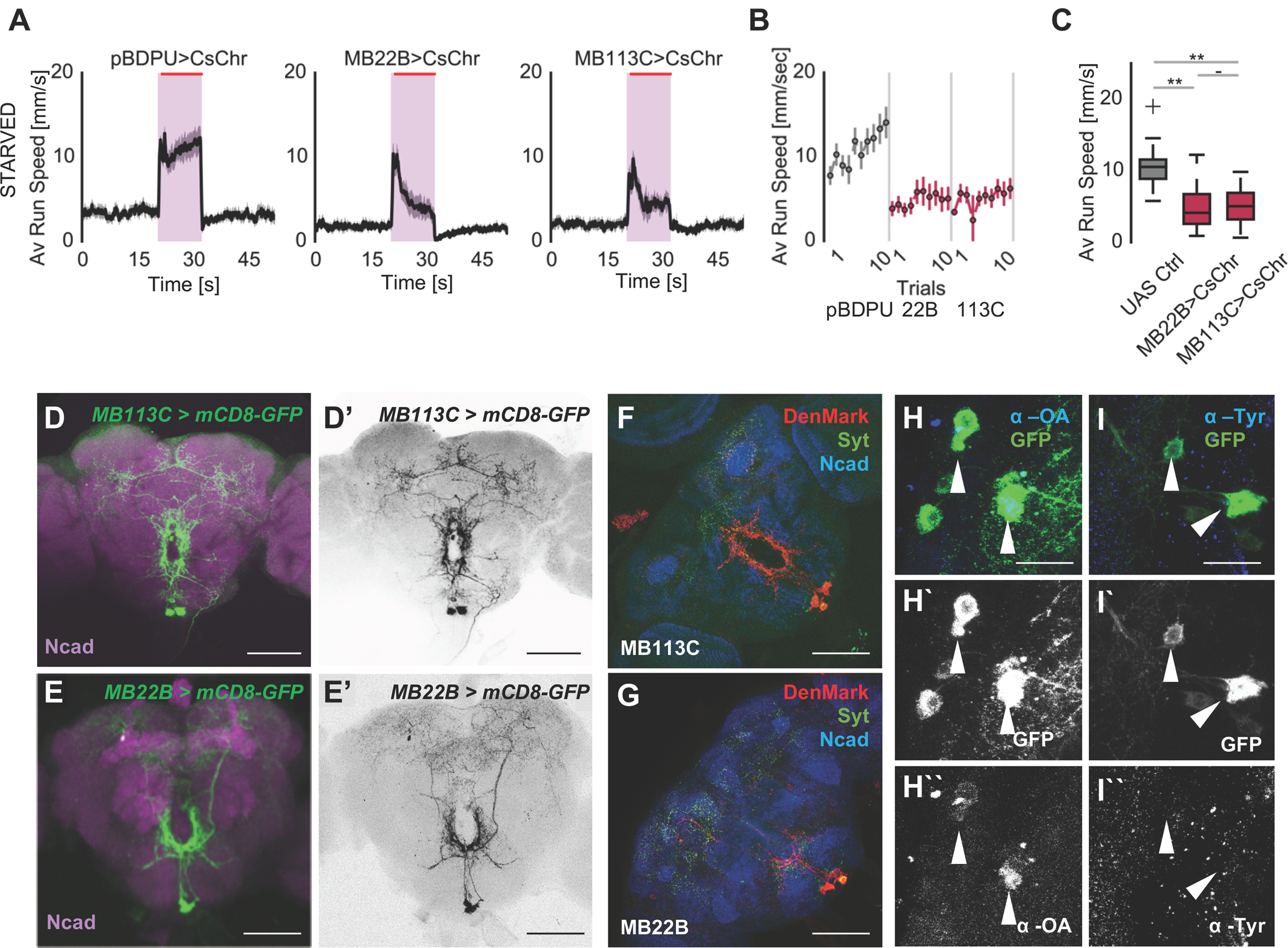
A subset of octopaminergic neurons called VPMs inhibit odor tracking. (A-C) Acute optogenetic of VPM neurons. MB22B harbors VPM3 and VPM4 neurons, whereas MB113C labels only VPM4 (Control: *pBDP-Gal4U>UAS-CsChrimson).* Acute manipulation of VPM neurons also prevented flies from tracking the odor. Average running speeds for *MB22B>UAS-CsChrimson* and *MB113C>UAS-CsChrimson* flies during odor exposure over trials (A). Average running speeds for *MB22B>UAS-CsChrimson* and *MB113C>UAS-CsChrimson* flies during odor exposure (C, n=10/10/10). (D-G) Expression patterns and polarity of MB22B and MB113C split-Gal4 lines *(UAS-mCD8-GFP* for expression and *UAS-DenMark* / *UAS-syt-GFP* for polarity analyses). VPM3 and VPM4 neurons have their dendrites in the SEZ (subesophageal zone) and PEZ (periesophageal zone) and project dense arborizations in various regions in protocerebrum. (H-I``) VPM4 neurons *(MB113C>mCD8GFP)* are indeed octopaminergic and express octopamine. MB113C labeling co-localizes with α-OA in the soma, but not with α-tyramine (Tyr). For all analyses, one-way ANOVA with Tukey’s HSD post hoc analysis was used. : ‘ - ’ > 0.05, ’ * ‘ p < 0.05, ‘ ** ’ p < 0.01, ‘ *** ‘ p < 0.001).

Anatomically, VPM 3 and 4 neurons extend dendrites within the sub- and periesophageal zones, and send projections to the MB and other higher brain regions (Fig. 6D-G). Line MB113C and MB22B overlap only in the neuron type, VPM4, indicating that this neuron was central to the observed behavior (Aso et al., 2014a). Antibody staining against octopamine confirmed the categorization as OAN (Fig. 6H-I``).

Hence, VPM4 neuron activity can regulate odor tracking - conceivably, and as suggested by the imaging data (see Fig. 5), by mimicking food reward, or in other words, reaching of the goal.

### VPM4 antagonizes MBON-induced food odor seeking

VPM4 sends axons into the MB area. Interestingly, at the level of the MB lobes, the innervation pattern of the MBON MVP2 overlaps partially with VPM4 in the γ1 MB lobe indicating that VPM4 and MVP2 might be directly connected (Fig. S7.1A-A3). Moreover, recent analysis of the MB connectome in the *Drosophila* larva revealed an octopaminergic neuron, named OAN-γ1, forming direct synapses onto MBON-γ1/γ2 neurons, which are proposed to mediate feed-forward inhibition similar to MVP2 (Eichler et al., 2017).

We investigated the possibility of similar connectivity using a recently published EM volume of an entire female adult *Drosophila* brain (Zheng et al., 2017). Reconstruction of MBON MVP2 and subsequent sampling of its synaptic inputs to the γ1 compartment quickly led to the identification of two putatively aminergic neurons (based on the presence of dense core vesicles). Further reconstructions identified these neurons morphologically to be VPM3 and VPM4 (Fig. 7A_1_-A_3_). Both neurons form numerous synaptic contacts onto MVP2’s dendrites exclusively in the γ1 compartment with VPM4 making about 50% more synapses than VPM3 (Fig. 7A_3_, Fig. S7.1B). VPM3/4 > MVP2 presynaptic boutons featured small clear core vesicles surrounding the active zone as well as parasynaptic large dense core vesicles suggesting both synaptic and non-synaptic transmission (Fig. S7.1C1,C2). Whether both types of vesicles release octopamine and/or an additional transmitter is not known. In addition, MVP2 also made a smaller number of reciprocal synaptic contacts back to VPM3 but not to VPM4 (Fig. S7.1B). We found no evidence that VPMs contact MVP2 in other regions of the MB lobes, although we cannot completely rule this out.

**Fig. 7.**
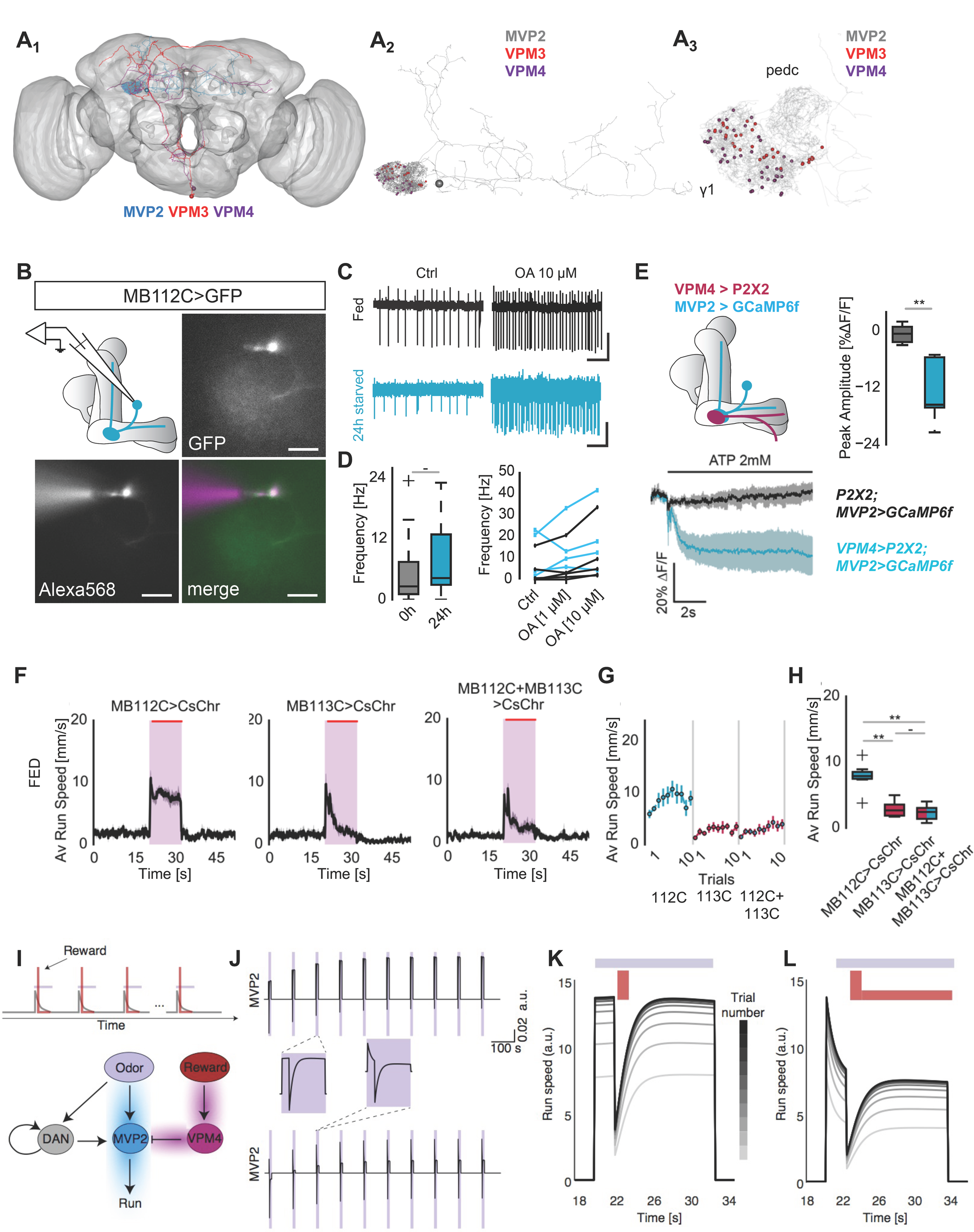
VPM4 modulates MVP2-dependent tracking. (A_1-3_) EM reconstruction reveals synaptic connections between MVP2 and VPM3 and 4. (A1) Skeletons of EM reconstruction of MVP2 (blue), VPM3 (red) and VPM4 (purple) on the neuropil of a whole fly brain. (A2) Red (VPM3) and purple (VPM4) indicate the synapses between VPMs and MVP2, respectively. (A3) Higher magnification of A2. Note that all VPM3/4 > MVP2 synapses are found in the γ1 lobe region of the MB. (B-E) Effect of OA application on MVP2 neurons firing rate. (B) Image of a MVP2 neuron visualized by *MB112C>mCD8-GFP* (green) and dye-filled with Alexa568 in a whole cell configuration (magenta; scale bar 20 μm). (C) Representative current traces of MVP2 cell attached recordings from fed and 24h starved flies, before and after bath application of OA (10 μM; scale bars 1 s, 20 pA and 4 pA). (D) Left panel: Average firing rates during 2 minutes of recording (n=11/7; unpaired T-test). Right panel: Effect of bath application of OA (n=5/4; fed vs. starved ns; OA concentration p<0.05; interaction ns; two-way repeated measures ANOVA). (E) Upper left panel: Scheme showing VPM4 and MPV2 neurons at the level of the mushroom body and the genetic combination of transgenes expressed in the fly used for the experiment. Blue: MVP2 expressed GCaMP6f *(MB112C-Gal4;UAS-GCaMP6f),* purple: VMP4 expressed P2X2 *(GMR95A10-lexA;lexAop-P2X2).* Lower left panel: Average traces ±SEM displaying % ΔF/F GCaMP fluorescence in MVP2 (lobe area) upon ATP application on brain in an *in vivo* preparation. Upper right panel: Box plots display peak amplitude of % ΔF/F GCaMP fluorescence in MVP2 upon ATP application. Turquoise: test group: *MB112C>GCaMP6f;VPM4-P2X2,* dark grey: control group: *MB112C>GCaMP6f;+/lexAop-P2X2.* n=5, T-test: 0.0066. (F-H) Epistasis experiment for VPM4 and MVP2 suggesting that VPM4 suppresses MVP2 induced odor tracking. In fed flies, simultaneous activation of MVP2 and VPM4 resulted in sustained reduction of odor tracking speed upon vinegar stimulation (n=7/7/8). (I-L) Model captures the effects of reward delivery. (I) We modeled two different types of reward, one transient (see also K) and a second transient and sustained (see also L) (red bars) to mimic the effects of Gr5a (peripheral sugar sensors) and Gr43a (peripheral and internal sugar sensors) activation (Fig. 4A), respectively. New circuit diagram includes VPM4 neurons that convey the reward to MVP2 (magenta shading). (J) Top: Gr5a activation is captured by applying a strong and transient reward. This reward transiently and strongly decreases MVP2 activity, while the continued input from the DANs enables a fast recovery. Bottom: Gr43a activation is captured by applying a weak and sustained reward following the strong and transient reward signal. This reward delays MVP2 recovery despite continued input from the DANs. (K) Applying a strong and transient reward in the model generates speeds which, following a transient decrease, recover to initial values, similar to the activation of peripheral sugar receptors, Gr5a (Fig. 4A). (L) Applying a weak and sustained reward (in addition to the transient) in the model generates speeds which, following a transient decrease, remain low throughout the trial, similar to the activation of internal sugar receptors, Gr43a (Fig. 4A). Model parameters as in Fig. 1, additionally with *R =* 1000, *r =* 0 in J (top), K and *R =* 1000,*r* = 0.05 in J (bottom), L. For all behavioral analyses, one-way ANOVA with Tukey’s HSD post hoc analysis was used. : ‘ - ’ > 0.05, ’ * ‘ p < 0.05, ‘ ** ’ p < 0.01, ‘ *** ‘ p < 0.001.

We used *ex vivo* cell attached recordings to test whether MVP2 neurons expressed receptors sensitive to OA released by VPM3/4. We found that on average the MVP2 neuron responded with an increase in spiking activity to 1 and 10 μM of OA stimulation strongly suggesting that these neurons express OA receptors (Fig. 7B-D). This was true for brains of starved and fed animals (Fig. 7D). In one brain of a starved fly, we observed the inhibition of spontaneous spiking possibly indicating different OA receptors or the presence of an inhibitory feedback loop with might involve additional neurons (Fig. 7D). In addition, we observed that some MVP2 neurons showed a high baseline activity while others remained relatively silent (Fig. 7D), perhaps indicating a state-dependent difference in baseline. These observations show that MVP2 neurons are indeed sensitive to OA. Next, we expressed the ATP-sensitive mammalian channel P2X2 (Yao et al., 2012) in VPM4 to directly test the effect of activating this OAN on MVP2 (Fig. 7E). As a proxy of neuronal activity, we monitored GCaMP6f fluorescence in MVP2 upon adding ATP to an *in vivo brain* preparation (Fig. 7E). Addition of ATP led to a significant decrease in GCaMP fluorescence in MVP2 neurons in the presence of P2X2 in VPM4 but not in controls strongly suggesting the existence of an inhibitory connection between these two neurons (Fig. 7E).

To test the role of the observed synaptic connection between VPM4 and MVP2 during goal seeking behavior, we argued that based on our observation that activation of VPM4 and inactivation of MVP2 showed similar phenotypes (i.e. reduction of odor tracking), activation of VPM4 could potentially override the activation of MVP2 and inhibit induced running after the odorant in the fed animal. To this end, we combined *MB112C-* and *MB113C-Gal4* with *UAS-CsChrimson* and applied the same protocol as above. Activation of only MVP2 led to significant odor tracking in response to vinegar in the fed fly as expected (Fig. 7F-H, Fig. S7.1D,E). Concurrent activation of both neurons (VPM4 and MVP2) showed that VPM4 activation with either Split-Gal4 line, MB113C and MB22B, was sufficient to override the activation of MVP2 (Fig. 7F-H, Fig. S7.1D,E).

In summary, MVP2 MBONs express OA receptors and can be inhibited by VPM4 activation. Activation of OA-VPM4 neurons is sufficient to suppress behavioral output mediated by the activation of MVP2. Taken together, we propose that a specific OA neuron, VPM4, receives external and internal sugar neuron input, which it transmits to MBONs, such as MVP2, to suppress persistent food odor tracking, once food has been found and ingestion of a nutritive food source has started.

### Modeling the role of MVP2 in integrating the lack or the finding of an expected reward

We included two types of reward in the existing model to investigate the effects of activating nutrient sensing neurons (i.e. Gr5a and Gr43a) and the action of VPM4 on MVP2 output (Fig. 7I, magenta shading). Now, both the activity of DANs and the reward from VPM4 modulate the output of MVP2 (Fig. 7I).

The first type of reward was modeled as a strong, but non-lasting input that mimicked the transient reward conveyed by the taste of sugar through for instance Gr5a neurons. Since the perceived reward due to the lack of ingestion and post-ingestion signals was short-lasting, we observed that MVP2 dynamics rapidly recovered during the odor, which continued predicting a possibly ‘better’ reward (Fig. 7J, top). This rapid recovery suggested that the synaptic time constant of MVP2 must be much faster than the time constant accumulating the persistence signal by the DANs. We were unable to infer the exact MVP2 time constant from the fitting since we fitted the behavioral data without reward; however, exploring different time constants did not change our prediction that the DAN time constant is longer than the inter-stimulus interval, suggesting that the model output was not sensitive to the exact value of this parameter (Table 1, see Methods). Transforming MVP2 output into running speed using the same nonlinearity as before (see Fig. 3J) generated speeds which decreased drastically when the reward signal was transmitted, but rapidly recovered to their initial value due to the continued input from the DANs (Fig. 7K; compare to optogenetic Gr5a activation in Fig. 4A).

For the second type of reward, we added a weak and constant component to strong transient signal from before, to obtain a sustained reward expected due to longer-lasting post-ingestion signals such as activation of internal sugar sensors (e.g. Gr43a) (Fig. 7I). This is in agreement with the intracellular recordings in OA neurons in honeybees mentioned above, which have shown a transient burst of high frequency firing followed by prolonged activation upon sugar feeding (Hammer, 1993). The presence of this sustained signal following the transient one prevented the recovery of MVP2 activity in the model despite continued input from the DANs (Fig. 7J, bottom). The running speeds remained low during odor presentation (Fig. 7L; compare to Gr43a activation in Fig. 4A). Therefore, our model can simultaneously capture the accumulation of negative experience due to lack of reward at the level of DANs, and the signaling of reward by VPM4. The reward signal acts to overwrite the DAN-mediated persistence as this dual information is integrated in the MBON MVP2.

Taken together, theoretical modeling supports our experimental data and suggests that a network of a subset of DANs, a specific OAN, and a MBON can integrate external and internal sensory stimuli and intrinsic motivation to promote and control internal and behavioral state-dependent goal seeking behavior (Fig. 8).

**Fig. 8.**
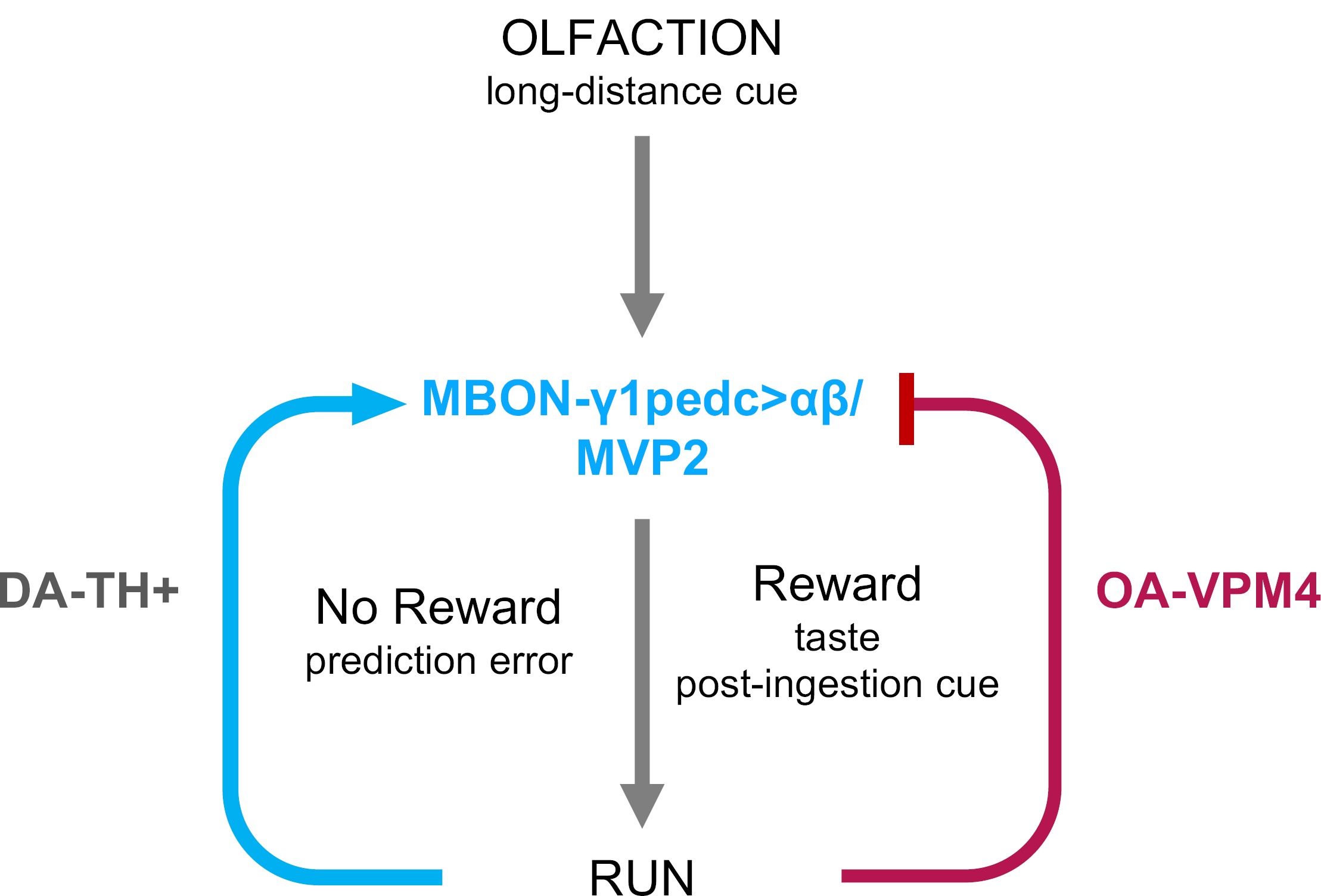
Schematic model. In the hungry animal, food odors induce a strong impulse to forage for food by following the odorant (Run). This tracking requires the MBON MVP2. In lieu of an expected reward, TH+ DANs, which are presumably activated through a combination of odor and high movement in lieu of reward, leads to reinforcement and higher perseverance. Once the animal encounters the desired reward, the food, it stops to evaluate and eat the food Post-ingestion signals such as internal sugar rise, will prevent the animal from leaving the food source and running again. Thus, taste as well as internal signals override olfactory stimulation and subsequent behavioral programs. This is mediated by the VPM4 OA neuron, which integrates hunger state and appetitive taste and counteracts the odor-induced, MVP2-dependent food foraging response. Of note, the exact nature and sign of the synapse between MVP2 and VPM is currently not known. We speculate that it is state-dependent and that VPM4 might release more than one neurotransmitter. Alternatively, there might be a negative feedback loop back to MVP2.

## Discussion

Theory predicts that any behavior an animal decides to engage in or withdraw from is a result of a cost-benefit calculation, where both cost and benefit strongly depend on the state and need of the subject (Schultz, 2015; Tymula and Plassmann, 2016). Similarly, an expected benefit that does not materialize, a so-called reward prediction error, rapidly decreases the animal’s interest in investing into getting this reward (Schultz, 2015). Here, we have shown that a hungry fly in lieu of an expected reward does not give up, but instead perseveres. In other words, this reward prediction error does not reduce but stimulate the very behavior that was previously unsuccessful. Furthermore, the observed behavior does not appear to require classical learning mechanisms; neither does it seem to induce long-term memory. Instead, we propose that the present paradigm uncovers a different role for the two important neuromodulators dopamine and octopamine, and shows that they are used to flexibly translate the animal’s ongoing experience into the most advantageous and appropriate, immediate behavioral response.

### Dopamine as a signal for perseverance

Mechanisms of being persistent in one behavior over another have previously been analyzed in flies. For instance, courtship of fly males and copulation with a female are maintained by dopaminergic neurons in the ventral nerve cord, where they counteract GABAergic neurons (Crickmore and Vosshall, 2013). In that scenario, DANs in the ventral nerve cord reinforce an ongoing behavior and prevent that the male disengages prematurely before successful insemination.

Why does the fly not give up running after a food cue although its behavior is not rewarded? The advantage of preserving energy by not following the odor cue would be to have sufficient reserves to follow a next, and hopefully this time rewarding, stimulus. Nevertheless, the hungrier the fly is, the higher its perseverance. Conversely, the fed animal, although it shows an initial odor reaction and as our imaging data suggests, detects the odor not different from the starved fly, makes little effort to follow the cue. It is evident that internal state-dependent motivation gates this behavior. In line with this, a sugar reward - we applied it using optogenetic stimulation of peripheral and internal sugar receptor neurons - instantaneously interrupted food seeking by odor tracking, so that the animal could engage into feeding. However, it did not enhance the appeal of the odor in future trials.

Our data therefore suggest that both the negative experience of a reward prediction error and the positive experience of the reward either do not induce long-term behavioral changes or such long-term behavioral changes are not opportune in the present task, but might be expressed under different or future circumstances. It is not obvious why it might be advantageous for a hungry fly, to not follow a food cue in the future, even though a past one was not rewarded. However, this might apply specifically for *bone fide* or frequently experienced food odors, while for neutral or more ambiguous odors learning - and presumably also unlearning through a reward prediction error - would occur (see also (Felsenberg et al., 2017)).

### Circuit mechanisms and the role of the mushroom body network

We have shown that the MBON MVP2/MBON-γ1>αβ is critical to promote efficient odor tracking. Moreover, it is also required for the choice of vinegar over humidified air in a simple choice assay, the T-maze. Consistently, MVP2 was previously shown to respond strongly to food odors including vinegar (Hige et al., 2015b). Interestingly, in the context of appetitive olfactory memory, MVP2 acts as a gate keeper during the expression of such memory, where it functionally counteracts the activity of another MBON to allow learned odor attraction (M4/6 or MBON-β`2) (Perisse et al., 2016). Conversely, MBON-β`2 mediates odor aversion and its activity is repressed by a vinegar odor-sensitive PAM DAN in the context of conflicting sensory information (Lewis et al., 2015; Owald et al., 2015). In these scenarios, MB KCs play an essential role. This appeared not to be the case in the present study. Inactivation of all KCs or just γ KCs had no impact on food odor tracking. By contrast, DANs play a key role as mediators of negative experience and promoters of perseverance, as further corroborated by our theoretical model. DANs, which are highly sensitive to odor, taste, post-ingestion signals and punishment such as physical pain (Huetteroth et al., 2015; Mao and Davis, 2009; Musso et al., 2015; Riemensperger et al., 2005), serve as teaching signals and imprint these negative and positive experiences into the MB network for instance by modifying the KC-MBON synapse (Cohn et al., 2015; Hige et al., 2015a; Owald et al., 2015). Interestingly, DAN activity correlates with motor activity and metabolic state of the animal (Berry et al., 2015; Cohn et al., 2015; Krashes et al., 2009; Placais and Preat, 2013). In particular, the activity of one PPL1 neuron, MV1, but not of another, V1, is higher in flies moving on a spherical treadmill (Berry et al., 2015). On the other hand, MP1 integrates and transmits hunger state to the mushroom body and thereby governs metabolic state-dependent appetitive memory expression (Krashes et al., 2009). In addition, optogenetic activation of sweet-taste coding tarsal projection neurons inhibits basal calcium responses in several PPL1 types, including MV1, MP1, and V1 (Kim et al., 2017). These previous data are consistent with our model that some DAN activity accumulates in the absence of an expected food reward and decreases in the presence of taste or food.

How do DANs modulate behavior without KC involvement? And where does the odor information come from if not through KCs? Connectomics recently showed that DANs synapse directly onto MBONs providing an anatomical substrate for a KC-independent role of DANs and MBONs (Eichler et al., 2017; Takemura et al., 2017). Furthermore, output neurons of the lateral horn project toward the MB and might transmit odor information by connecting to MBONs or DANs (Aso et al., 2014a). Theoretical modeling based on our experimental findings supports that a direct synaptic connection between DAN and MBON is sufficient to explain the behavioral data. DANs, which respond to external and internal cues including hunger, odor and movement (Berry et al., 2015; Lewis et al., 2015; Mao and Davis, 2009; Placais and Preat, 2013; Riemensperger et al., 2005), represent the ideal candidates to reinforce MVP2-mediated motor behavior during repeated odor presentation in the absence of reward; they modulate the output of MVP2, effectively decreasing its activation threshold and driving the animal to run faster and longer in later trials. Notably, such a direct modulation without the involvement of KCs might be the key as to why no long-term memory is observed or required for perseverance in food odor tracking. It is possible that this reinforcement between DANs and MVP2 occurs through a direct feedback loop (Ichinose et al., 2015; Zhao et al., 2018). Zhao et al. showed that a feedback loop between an MBON and a DAN allowed neuronal activity to persist, which in their case was important to maintain a courtship memory. Our modeling results demonstrated that a key ingredient for the observed persistence in behavior is the long time constant operating during repeated odor presentations and the accumulation of a reward prediction error, consistent with a direct feedback loop from recurrent circuit interactions. Future experiments will address whether such a putative direct feedback loop between MVP2 and DANs indeed induces accumulation of activity in the DANs.

What is the identity of the DAN signaling the error and promoting perseverance? An obvious candidate is the MP1 or PPL1-γ1pedc DAN, which overlaps with MVP2 in its innervation of the MB (Aso et al., 2014a). Moreover, the negatively-reinforcing MP1 DANs prevent the expression of appetitive memory in fed flies through a hunger state-dependent peptidergic mechanism (Krashes et al., 2009; Perisse et al., 2016). Finally, a recent study suggested that MP1 and MVP2 are required for the aversion of an odor associated to the repeated defeat in fights between male flies (Kim et al., 2018). Nevertheless, *shi^ts1^-* mediated inactivation of the MP1 DAN neurons had no effect on perseverant odor tracking of the hungry animal (data not shown), suggesting the involvement of more or different, yet to be identified, DANs.

Similar to dopamine, OA has been implicated in appetitive olfactory learning and memory, where it gates memory expression in a hunger-dependent manner upstream of OA receptor-expressing DANs (Burke et al., 2012). Activation of Tdc2+ neurons effectively replaced sugar reward during learning and reliably induced short-term, but not long-term, appetitive memory. Interestingly, not only PAM DANs, but also OA receptor-expressing PPL1 DANs, which might be innervated by VPMs, were implicated in appetitive memory formation in this study (i.e. PPL1 MP1). However, activation of VPMs 3 and 4 alone did not induce learning (Burke et al., 2012), indicating that they are either not involved in this interaction or that essential additional signals are missing. Our anatomical data show that VPM3/4 synapses directly on the MVP2 MBON again suggesting that the identified mechanism bypasses the KCs but still modulates MBONs. This is also supported by our imaging data showing that activation of VPM4 inhibits MVP2. Through this mechanism, MBONs would receive the combined input of sensory (internal and external reward cues), present (online feedback via DANs) and past (lasting experience via DANs and KCs) information to control a behavioral action in the best interest of the animal. The model corroborates the influence of this (positive and negative) combined input: different types of reward can powerfully modulate the behavior induced by accumulation of negative experience by DANs, through specific OANs by impacting MBON output transiently or lastingly. While we start to understand how short- and long-term olfactory memories are formed, this single fly assay reveals an additional function of the MB network in behavioral perseverance and withdrawal, and provides a powerful way to study the interaction between octopamine and dopamine in the context of present experience.

### Food search and feeding - Sequential and antagonistic behaviors

In addition to insights into goal seeking and reward prediction mechanisms, our data exposes interesting aspects of feeding related behavior. Behavior is typically expressed as a sequence of antagonistic behaviors. Seeds et al. have shown this beautifully on the example of grooming behavior (Seeds et al., 2014). Multiple studies have analyzed the neuronal mechanisms underlying attraction to food and food seeking, food finding, and feeding itself in *Drosophila* and other animals (Thoma et al., 2017). Nevertheless, how these behaviors that are typically mutually exclusive and consecutive, relate and regulate each other is not well understood as prior work has focused primarily on long-term effects of hunger and feeding (Pool and Scott, 2014; Su and Wang, 2014). Finding food should induce food evaluation and feeding, while suppressing food search. Satiety counteracts the drive to find food, but a too small or not palatable food source should quickly re-induce food search, as it can be considered a reward prediction error. Similarly, feeding itself represents a series of consecutive behaviors that are, in part, mutually exclusive, and suppress or are suppressed by locomotion (Mann et al., 2013; Thoma et al., 2016; Thoma et al., 2017). Moreover, ingestion is regulated by feeding through a group of cholinergic interneurons that receive pharyngeal taste information (i.e. through Gr43a neurons), which maintains ingestion behavior (LeDue et al., 2015; Yapici et al., 2016).

Although a number of behaviors are easily distinguishable into meaningful substructures (e.g., courtship, larval navigation, grooming), foraging is rarely studied in its entirety. The use of a single animal olfactory treadmill has allowed us to dissect different aspects of food search and discovery. In particular, how does food discovery suppress food search, if the sensory cue, the odor, is still present and as our data suggests still forwarded to higher brain centers? We have argued that candidate mechanisms include a neuron or a pathway of neurons capable of integrating taste and hunger state and projecting it to neurons processing long-distance food cues such as odors that are critical for food search itself. The OAN VPM4 fulfills these requirements. Its dendrites in the SEZ are sensitive to taste in a hunger state-dependent manner, and it connects directly to the MBON MVP2 that is required for food odor tracking. Importantly, activation of VPM4 inhibits MVP2 activity and MVP2-induced odor tracking strongly suggesting an inhibitory connection between VPM4 and MVP2. Finally, our modeling data support the simple scenario we propose and show that a minimal network of the three identified types of neurons (i.e. VPM4, MVP2, DANs) can produce the behaviors that we have observed in our experiments.

### A specific role for octopamine/norepinephrine in taste-modulated behaviors?

Here, we pinpoint a role for a specific OAN in taste and feeding in flies. In mammals, the role of NE signaling has been analyzed primarily in the context of the brainstem LC. Notably, the LC also contains DANs, which play a role in memory consolidation (Takeuchi et al., 2016). Another structure containing a large number of NE neurons is the NST. Interestingly, the NST is also part of what is thought of as the central gustatory pathway (Smith and Lemon, 2007). The rostral part of the NST receives input from the tongue as well as input from the vagus nerve, and therefore integrates external and internal information (Smith and Lemon, 2007). In addition, it serves as an area of multisensory integration of taste with temperature, texture, and odor (Escanilla et al., 2015; Wilson and Lemon, 2013). Neurons in the NST project to multiple brain regions including the amygdala, hypothalamus, and insular cortex (Carleton et al., 2010), all of which receive internal state as well as other sensory information. In fact, electrical stimulation of the NST led to the release of NE in the central amygdala (Myers and Rinaman, 2002) indicating that this activity might influence feeding decisions, which are also regulated by this brain structure (Douglass et al., 2017).

We hypothesize that the here identified elements of a circuit consisting of specific OANs, an MBON and DANs might play a fundamentally similar role as NE neuron-containing circuits in the mammalian brain stem; integration of internal and external context to organize behavior in a flexible and context-dependent manner.

## Acknowledgements

We would like to acknowledge Lasse Bräcker and Jörg Henninger for the development of an early version of the treadmill assay. We thank Christian Schmid for his help with the 4- arm olfactometer assay optimization. Furthermore, we are grateful to Stefan Precht and the Max-Planck-Institute of Neurobiology workshop for their excellent work. We are grateful to Vivek Jayaraman and Armin Bahl for help with the treadmill assay. We thank Anja B. Friedrich for technical help with histology and confocal imaging and Heidi Miller-Mommerskamp for help with general fly husbandry. We are very grateful to Yoshinori Aso and Gerald Rubin for sharing fly lines prior to publication. We also thank Matthieu Louis, Hiromu Tanimoto, Scott Waddell and members of the Grunwald Kadow lab for comments on the manuscript. We further thank Scott Waddell, R. Roberts, N. Massood-Panah and I. J. Alifor for their contributions to the EM tracing.

This work was funded by the Max-Planck-Society (to JG and IGK) and the Technical University of Munich (to JG and IGK), an ERC starting grant (637472, to IGK), the Marie-Curie-Training Network FliAct (to IGK), a Capes-Humboldt postdoctoral fellowship (to MW) and the German Research Foundation (CRC870 (A04), INST 95/1382-1 and INST 95/1419-1, to IGK). All EM reconstruction work was supported by the Howard Hughes Medical Institute (to DDB), the Wellcome Trust (203261/Z/16/Z), an ERC Consolidator grant (649111) and core support from the MRC (MC-U105188491) to GSXEJ.

## Author contributions

SS designed and built the spherical treadmill assay with help and input by IGK. JFDB designed and carried out all imaging and electrophysiology experiments with the exception of PN imaging, which was carried out KPS. MW and JG developed the theoretical models based on experimental data provided by SS. LPL designed and built the 4-arm maze and carried out the T-maze screen for MBON lines involved in vinegar attraction. LMF carried out and analyzed the 4-arm maze experiments. PS, AES, CBF, NF, SCS, SL, DB, MC, and GSXEJ carried out and analyzed all experiments relating the EM reconstruction. SS and IGK conceptualized the study with input by JFDB. IGK oversaw and directed the study and wrote the manuscript with contributions of all authors.

## Materials and Methods

### Fly Husbandry and Lines

Flies were raised at 25°C, 60% humidity, with a 12/12 hours light/dark cycle on a standard cornmeal media. For optogenetic experiments, adult flies were collected at eclosion, kept under blue light only conditions (470nm, 0,05 μW/mm2) on an all-trans-retinal supplemented food (1:250). Fly lines were obtained from the Bloomington stock centers or directly from Janelia Research Campus. Fly lines used in the study were obtained from the Bloomington Drosophila Stock Center: Tdc2-Gal4, GMR95A10-LexA, UAS-Shibire^ts1^, UAS-dTrpA1, UAS-CsChrimson, UAS-DenMark, UAS-syt-GFP, UAS-mCD8-GFP, LexAop2-mCD8-GFP, UAS-GCaMP6f, UAS-GCaMP3, lexAop-P2X2 and GH146-Gal4. All Split-Gal4 lines including MB22b, MB112c, and MB113c were obtained from the Janelia Research Campus Flylight collection.

### Spherical Treadmill behavioral assay

The spherical treadmill was built according to (Seelig et al., 2010) with several modifications to accommodate olfactory instead of visual stimulation protocols. Fly tethering was performed under cold anesthesia and flies were immediately transferred onto the treadmill. After 3 min of acclimatization, experiments were initialized and controlled via a custom-written Python program. Flies that failed to acclimatize and reach a minimum speed of 1.5 mm/s before the first stimulus were discarded. An experiment consisted of 10 consecutive trials (with the exception of 6 trials for CO_2_ experiments to prevent anesthesia), which were separated by semi-randomized intervals of 60 ± 2-20 s. Each trial was recorded for a minimum of 52 s. The recording was divided into prestimulation (20 s), stimulation (dependent on experimental procedure) and poststimulation (30 s) periods. For open-loop experiments, the stimulation period was 12 s. The closed-loop experiments utilized a short open-loop phase (2 s), followed by a closed-loop phase. In the closed-loop phase, the fly controlled the odor stimulation length, i.e., the odor channel was kept open as long as the online speed criteria were met (>0 mm/s for 100 ms). The online speed data acquisition rate in the cardinal directions (yaw, pitch and roll) was ~4kHz for all experiments. The recorded speed data were down-sampled to 10 Hz by summation. Butterworth filtering was employed in 2D locomotion trajectory reconstruction. All data analyses were performed with Python 2.7, numpy 1.8, scipy.stats (0.14) and pyvttbl (0.5.2.2). Running and absolute turning average speeds were calculated as averages of 100 ms data points collected at cardinal directions in the respective phases of a trial. To minimize the impact of tethering artifacts, 100 ms data points for absolute turning speeds of each single trial were filtered with average absolute turning values, which were computed from the whole length of a respective trial. Average run time was defined as the initial uninterrupted running (speed > 0 mm/s) bout time length upon odor contact. Average run activity was measured as the fraction of odor stimulation time, where flies showed running speed higher than 0 mm/s. A stop was defined as not moving (0 mm/s) for at least 100 ms. Data visualization was done with matplotlib (1.4.2). Optogenetic activation on the ball was achieved by using a single high-power mounted LED at 617 nm, calibrated at 30 W/mm^2^ (M617, Thorlabs). Light stimulation in the absence of frontal air or odor stimulation induced some attraction and forward running toward the light source. This attraction was independent of the genotype of the animal and was seen also in wildtype Canton S flies (Fig. S3.2H). Similarly, simultaneous odor and light stimulation had not effect on Gal4 control flies (Fig. S7.2)

For appetitive olfactory stimulus delivery on the treadmill, a custom-made PTFE (Teflon) 4 mm tube was used and stationed at 3 mm distance from the tethered fly. The air speed was set to 100 ml/min via a Natec Sensors mass-flow controller. A balsamic vinegar solution (Alnatura Aceto Balsamico, Germany) was prepared daily at 20% v/v dilution in 100 ml Schott bottles. The vinegar concentration was measured with a miniPID (Aurora Scientific, miniPID 200B) and Arduino Uno at 100 Hz. PID recordings were filtered with Butterworth. The miniPID was calibrated with ethyl-butyrate according to (Semmelhack and Wang, 2009). For CO_2_ delivery, a CO_2_ stream was injected via a PTFE syringe inserted into the 100 ml/min pressured air main stream. The auxiliary line carried pure CO2 at 50 ml/min.

### Optogenetic and olfactory 4-arm maze

The 4-arm arena is based on the optogenetics-only arena described in (Aso et al., 2014b). The air/odor delivery was achieved via 4 passive solvent channels (Schott bottles containing Millipore water or vinegar solution 20% v/v) and a rotary pump (Thomas G12/01-4 EB). The rotary pump (~200 ml/min) was connected to an outlet at the arena center. The negative pressure generated by the pump facilitates drawing headspace in the passive solvent channels. For rapid switching between odor channels and target quadrants, a set of solenoid valves (Festo MFH-3-MF) were used. For optogenetics, a custom assembled LED array (Amber SMD PLCC2) was utilized to stimulate each quadrant of the arena at 617nm. The arena was illuminated via IR-LEDs and experiments were recorded with a CMOS camera (FLIR Flea3 MP Mono). The behavioral analysis expressed as preference index ((number of flies in stimulus quadrants - number of flies in non-stimulus quadrants) / total number of flies). Hardware control and data acquisition was achieved via Arduino Mega and in-house MATLAB scripts.

### T-maze

The two-choice population assay or T-maze was performed as previously described in (Lewis et al., 2015). Briefly, flies were tested in groups of ~60 in a non-aspirated T-maze and were allowed 1 minute to respond to stimuli. Experimentation was carried out in climate-controlled boxes at either 22-25°C or 32°C and 60% rH. A preference index (PI) was calculated by subtracting the number of flies on the air side by the number of flies on the stimulus side and normalizing by the total number of flies. Statistical analysis was performed using one-way ANOVA and the Bonferroni multiple comparisons post-hoc test using Prism GraphPad 6.

### Immunohistochemistry

Adult (4-7 days old) fly brains were dissected, fixed and stained as described previously (Lewis et al., 2015). All microscopy was performed at a Leica SP8 confocal microscope. Images were processed using ImageJ and Photoshop. The following antibodies were used for the neurotransmitter stainings: anti-OA (ABD-029, Jena Bioscience: mouse anti-conjugated octopamine (monoclonal) MAB OA-1) and anti-Tyr (AB124, Hemicon: rabbit anti-p-tyramine (polyclonal)), and mouse anti-ChAT (Yasuyama et al., 1995). The following antibodies were used to stain for (i) GFP: 3H9 primary antibody (specific to GFP1-10, monoclonal, Chromotek, 1:100), (ii) RFP: primary antibody 632496, Clontech: Living colors, rabbit anti-DsRed polyclonal (1:250), and (iii) Ncadherin: DSHB rat DN-EX #8 (1:100). Secondary antibodies: anti-rat Alexa568 (molecular probes, 1:250), antimouse Alexa488 and Alexa633 (molecular probes, 1:250), anti-rabbit Alexa568 and Alexa633 (molecular probes, 1:250).

### Electron microscopy and connectomic analysis

Reconstructions are based on an ssTEM (serial section transmission electron microscope) dataset comprising an entire adult fly brain (Zheng et al., 2017). Neuron skeletons were manually reconstructed using CATMAID (http://www.catmaid.org) (Saalfeld et al., 2009; Schneider-Mizell et al., 2016). MBON MVP2 was initially identified by sampling downstream of KCs in the γ1/peduncle compartment. VPM3 and VPM4 were found by semi-random sampling of synaptic inputs of MVP2 in the γ1 compartment. For identification, their microtubule-containing backbones were reconstructed and compared with published light-level data (Aso et al., 2014a; Busch et al., 2009). Subsequently, their axonal branches in the γ1 compartment were reconstructed to completion and synaptic sites were annotated. Synaptic connections described here represent fast, chemical synapses matching previously described typical criteria: thick black active zones, pre-(e.g. T-bar, vesicles) and postsynaptic membrane specializations (Prokop and Meinertzhagen, 2006). Visualisation and analysis were performed using open-source R (https://github.com/jefferis/nat and https://github.com/jefferis/elmr; (Manton et al., 2014)) and Python (https://github.com/schlegelp/pymaid) libraries.

### Electrophysiology

Cell attached patch-clamp recordings were performed on explant brains from 1 -3 days old fed and 24h starved *MB112C-Gal4;UAS-GFP* flies. Brains were dissected in ice-cold artificial adult hemolymph-like solution (AHL) containing (in mM): 103 NaCl, 26 NaHCO_3_, 3 KCl, 1 Na_2_HPO4, 4 MgCl_2_, 1.5 CaCl_2_, 5 TES, 10 D-glucose, 10 D-trehalose. After dissection, explant brains were transferred to a recording chamber, where they were maintained using a custom-made harp-shaped grid. The preparation was perfused during the entire experiment with AHL continuously bubbled with 95% O2/5% CO_2_ (~2mL/min). MVP2 neurons were identified using a Leica DM6000FS fluorescent microscope equipped with a 40x water immersion objective and a Leica DFC360 FX fluorescent camera. Prior to the recording, the glial sheath surrounding the MVP2 cell bodies was digested and removed using a pipette filled with 0.5 mg collagenase IV/ml in AHL. The exposed cell bodies were patched with 7-9 MΩ resistance patch pipettes filled with (in mM): 136 KMeSO_4_, 4 KCl, 0.022 CaCl_2_, 10 HEPES, 0.1 EGTA, 4 MgATP, 0.5 Na_2_GTP, 0.05 Alexa Fluor 568 Hydrazide. Recordings were acquired in cell attached mode using a Multiclamp 700B amplifier and a Digidata 1440A digital-analog converted driven by the Clampex10.3 software. Current traces were digitized at 20 kHz and on-line filtered at 3 kHz. Spontaneous firing activity was recorded for 2 minutes, 3 minutes after the formation of a gigaseal. Successive concentrations of OA (1 and 10 μM) were applied through the perfusion system for 10 minutes before recording. Data were analysed using the IgorPro6 software (WaveMetrics).

### Ex-vivo and in-vivo Calcium Imaging

Calcium imaging experiments were performed on 3-7 days old Gr43a-Gal4;UAS-GCaMP6f flies (for *ex vivo* experiments), 4-7 days old GH146-Gal4;UAS-GCaMP3 flies (for *in vivo,* odor stimulation), 4-8 days old Tdc2-Gal4;UAS-GCaMP6f flies (for *in vivo,* fructose stimulation), and on 4-7 days old R95A10-lexA,lexAop-P2X2;MB112C-Gal4;UAS-GCaMP6f flies and their controls +,lexAop-P2X2;MB112C-Gal4;UAS-GCaMP6f (for in vivo, ATP stimulation experiment). *Ex vivo* experiments were realized on explant brains prepared and maintained in the recording chamber as described above for electrophysiological recordings. These experiments were carried out in static conditions in 500 μL AHL. 50 μL fructose (1 M) was added with a pipette one minute after baseline acquisition to a final concentration of 100 mM in the bath. For experiments including stimulation of P2X2 with ATP, ATP was added to the buffer on top of the brain of a living fly to a final concentration of 2 mM. Preparations of flies for *in vivo* experiments were prepared as previously described (Bracker et al., 2013). A custom-made odor delivery system (Smartec, Martinsried) was used for vinegar stimulation. The odor was delivered in a continuous humidified airstream (1000 mL/min) through a 8-mm Teflon tube placed ~1 cm away from the fly. For taste stimulation, a syringe needle (MicroFil) was mounted on a micromanipulator (Narishige). A drop of fructose (1 M in distilled water) was delivered to touch the labellum during 10 s. Stimulus application was monitored by a stereomicroscope as described previously (Hussain et al., 2016).

For *in vivo* and *ex vivo* experiments, preparations were imaged using a Leica DM6000FS fluorescent microscope equipped with a 40x water immersion objective and a Leica DFC360 FX fluorescent camera. All images were acquired with the Leica LAS AF E6000 image acquisition software. *Ex vivo* data were acquired at a rate of 1 frame/s for 10 minutes, without binning. *In vivo* data were acquired at a rate of 20 frames/s for 75 s with 4x4 binning mode. Changes in fluorescence intensity were measured in manually drawn regions of interest (ROI) using the LAS AF E6000 Lite software. Relative changes in fluorescence intensity were defined as ΔF/F = 100* (F_i_ - F_0_)/F_0_ for the *i* frames after stimulation. Fluorescence background, F0, is the average fluorescence of 60 frames (1 min; *ex vivo*), 15 frames (750 ms; *in vivo* odor) or 20 frames (1 s; *in vivo* taste). Pseudocolored images were generated using a custom-written MATLAB program and ImageJ.

### Model

We assumed that the DAN and MVP2 activities at time *t* are governed by the following set of equations:

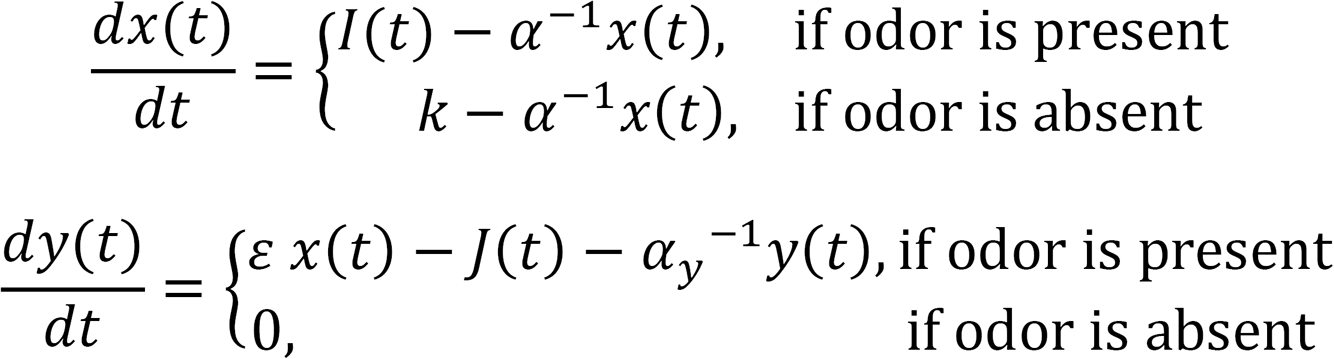

where *x(ť)* denotes DAN activity and *y(t)* denotes MVP2 activity. During inter-stimulus intervals, the odor input is set to zero, so MVP2 activity remains constant at a baseline. During odor presentation, the input is non-zero, so MVP2 is active and is modulated by the DANs. We modeled the input to the DANs *I(t)* as a decaying exponential of amplitude *A* and time constant *τ* (Fig. 3G). Projection neurons (see the transient traces in Fig. S3.3E) have connections with different populations of DANs, and we assume that the odor information is relayed to the DANs by the PNs. *k* is an auxiliary constant that sets the baseline in the absence of reward at each trial, *α* is the DAN time constant, *J(t)* is the reward (Fig. 7I), *α_y_* is the MVP2 synaptic time constant and *ε* is the time constant of the synapse between DANs and MVP2. We applied a nonlinearity to MVP2 output and scaled the result by a constant *S* to get speed values that have magnitudes in the same range as the data:

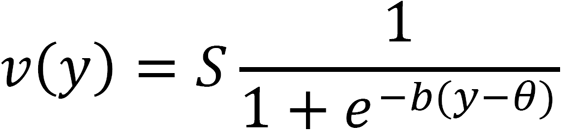

where *b* is the gain of the nonlinearity and *θ* its threshold. The reward in Fig. 7I is modeled as:

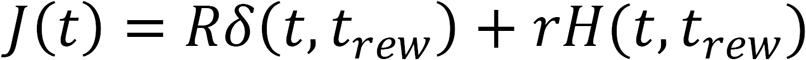

where *δ(t, t_rew_)* is the Kronecker delta function: *δ(t, t_rew_) = 1* if *t = t_rew_* and 0 otherwise. The Heaviside step function *H(t,t_rew_)* is equal to 1 if *t ≥ t*_rew_, and 0 otherwise. Both *R* and *r* are constants that modulate the strength of the reward, and for the transient reward (corresponding to Gr5a activation) we set *r =* 0.

### Model fitting and parameters

We used the data from Fig. 1E to fit our model’s parameters by minimizing the mean squared error between the model output (predicted running speed) and the data (measured running speed). As the data we used came from experiments without activation of reward, we set *R* = 0, *r = 0* in our initial model (Fig. 3G-K). We also set *ε = 1* without loss of generality. We fitted the DAN time constant *α,* the gain of the nonlinearity *b,* and the scale *S*. We fixed the time constant of MVP2 activity *α_y_* and the threshold of the nonlinearity *θ.* We chose the threshold values according to exploratory simulations where *θ* was fitted while other sets of parameters remained fixed. For the MVP2 time constant *α_y_,* we explored a range of values reported in Table 1, according to the criteria that it should be shorter than the period of odor presentation (12 s) to allow rapid MVP2 recovery after reward presentation during the odor. Although the exact values of the fitted DAN time constant, *α,* changed with *α*_y_, they remained longer than the inter-stimulus interval.

**Table 1:** Fitted model parameters after 150 Monte Carlo simulations with different initial conditions for DAN activity [0.04, 0.08] and different stimulus amplitudes [1.8, 2.3]. For each simulation, the optimization routine iterated until the convergence criteria of the error being <10^−5^ was achieved.

| Simulation set | Fitted DAN time constant ($\alpha$) | Fitted nonlinearity gain ($b$) | Fitted scale ($S$) |
| --- | --- | --- | --- |
| $\alpha_y = 1$, $\theta = 0.25$ | 162.6±13.0 | 9.5±2.2 | 50.8±20.7 |
| $\alpha_y = 2$, $\theta = 0.45$ | 177.3±9.8 | 4.3±0.3 | 36.2±4.2 |
| $\alpha_y = 5$, $\theta = 0.95$ | 197.3±5.2 | 1.6±0.1 | 27.7±0.6 |

### Trajectories

We randomly sampled 7 flies for each condition, and for these animals we selected the first, third, eighth, and last trials to show the trajectories. From the ball dynamics, we reconstructed the Cartesian coordinates of the fly in a flat surface using the transformations from (Seelig et al., 2010) and applied a Butterworth filter to smooth the paths.

---

**Fig. S1.1.**
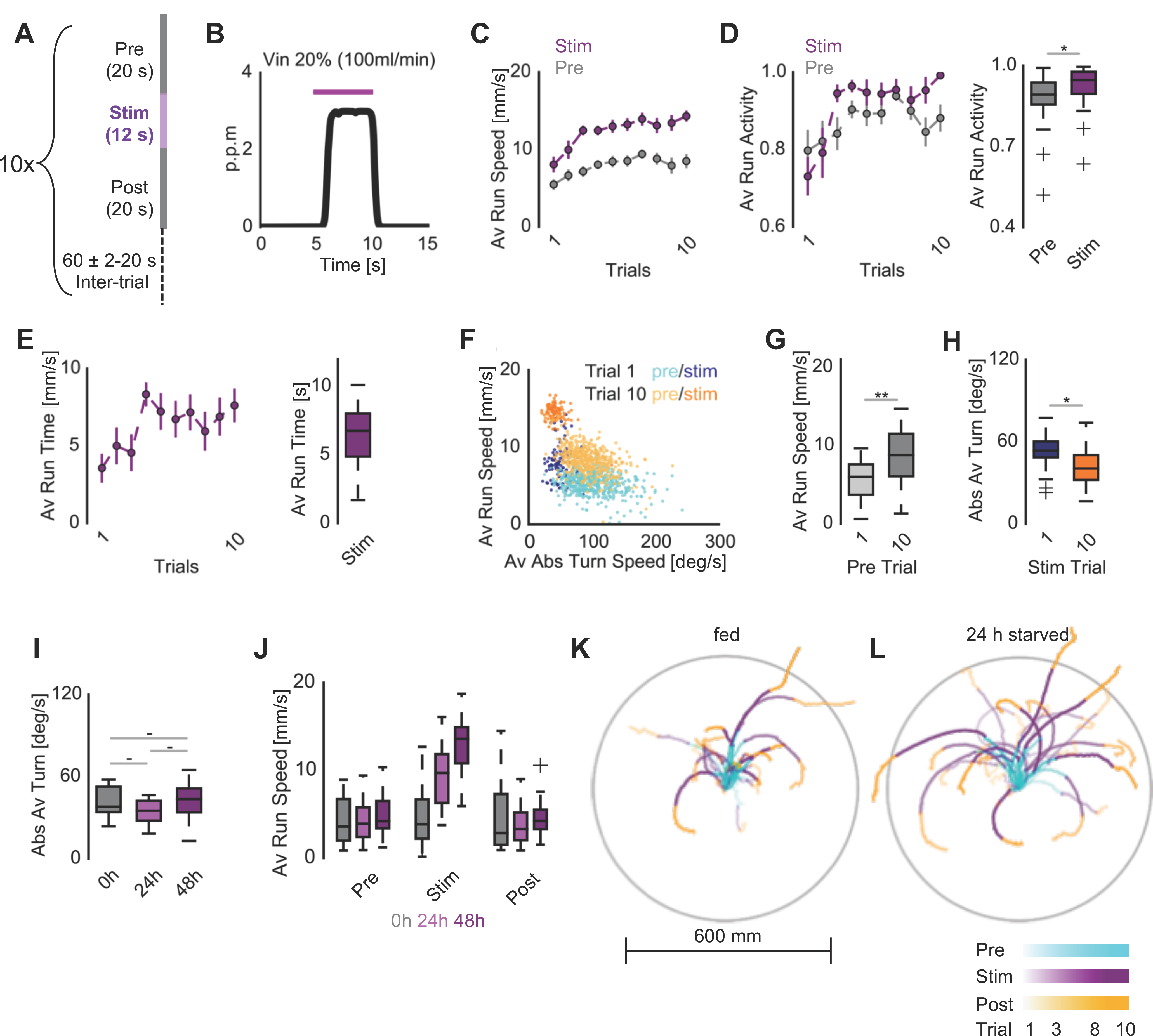
(A) Experimental protocol: Inter-stimulus periods were randomized in time (60 ± 2-20 s) to prevent odor onset predictability. The locomotor behavior was processed in two cardinal directions, runs and turns, at 10 Hz. This protocol was repeated 10 x per animal. (B) Representative vinegar concentration of a single application as used in treadmill behavioral experiments, measured with a photoionization detector (PID). After ~800ms latency at onset, odor concentration plateaued quickly and odor concentration stayed constant throughout the application. For visualization purposes, Butterworth filter was used for the representative PID trace. (C) Average running speed of 18 flies over trials with SEM. (D) Left: Average running activity over trials. Running activity is defined as a fraction of time where flies showed >0 mm/s running speeds for 100 ms over the defined odor pre-stimulation and stimulation periods. Flies were more active in running compared to the pre-stimulation periods (n=18, paired T-test). (E) Left: Average running lengths during the first bout of running upon start of vinegar exposure. Longer running bouts were observed over trials. Right: Average running bout length at the odor onset for all flies. (F) Evolution of the two-dimensional behavioral space on the spherical treadmill, constructed from 100 ms chunks of average running and absolute turning of 18 flies. Given the choice of run or turn, in average, flies executed turns in higher probability during in pre-stimulation period in the first trial (dark and light blue) than in the last trial (dark and light orange). The odor exposure shifted behavioral to longer runs at later trials from more turns at earlier trials. (G) Average running speed during pre-stimulation phases of trial 1 compared to trial 10. Speed increased during pre-stimulation phases with every trial reflecting the increase of speed during stimulation phases over trials (n=18, paired T-test). (H) Comparison of average absolute turning speed between trial 1 and 10. On average, flies turned less in the last trial compared to the first trial (n=18, paired T-test). (I) Average absolute turning speeds of hungry flies during vinegar exposure. Starvation did not change turning speed compared within any group (n=10/10/11, one-way ANOVA with Tukey’s HSD post hoc analysis). (J) Average running speed of flies during vinegar exposure in the closed-loop paradigm, including pre- and post-stimulation. Expanded from Fig 1L (n=20/18/19, filtered with Iglewicz and Hoaglin's robust test for multiple outliers). (K-L) 2D representation of tracks of 7 randomly chosen flies/condition for fed and 24 h starved animals. The lighter the shading of the trial, the earlier in the experiment. Trials 1,3,8, and 10 were plotted for all 7 flies. To facilitate graph interpretation a grey circle was drawn at 300 mm from the starting point of the fly. The absolute heading direction displayed is irrelevant as the odor was always kept in front of the animal. Note that hungry flies show significantly straighter and longer tracks as compared to fed flies, which do not move away much from their starting position. (n=7/7). For all analyses, statistical notations are as follows: ‘ - ’ > 0.05, ’ * ‘ p < 0.05, ‘ ** ’ p < 0.01, ‘ *** ‘ p < 0.001.

**Fig. S1.2.**
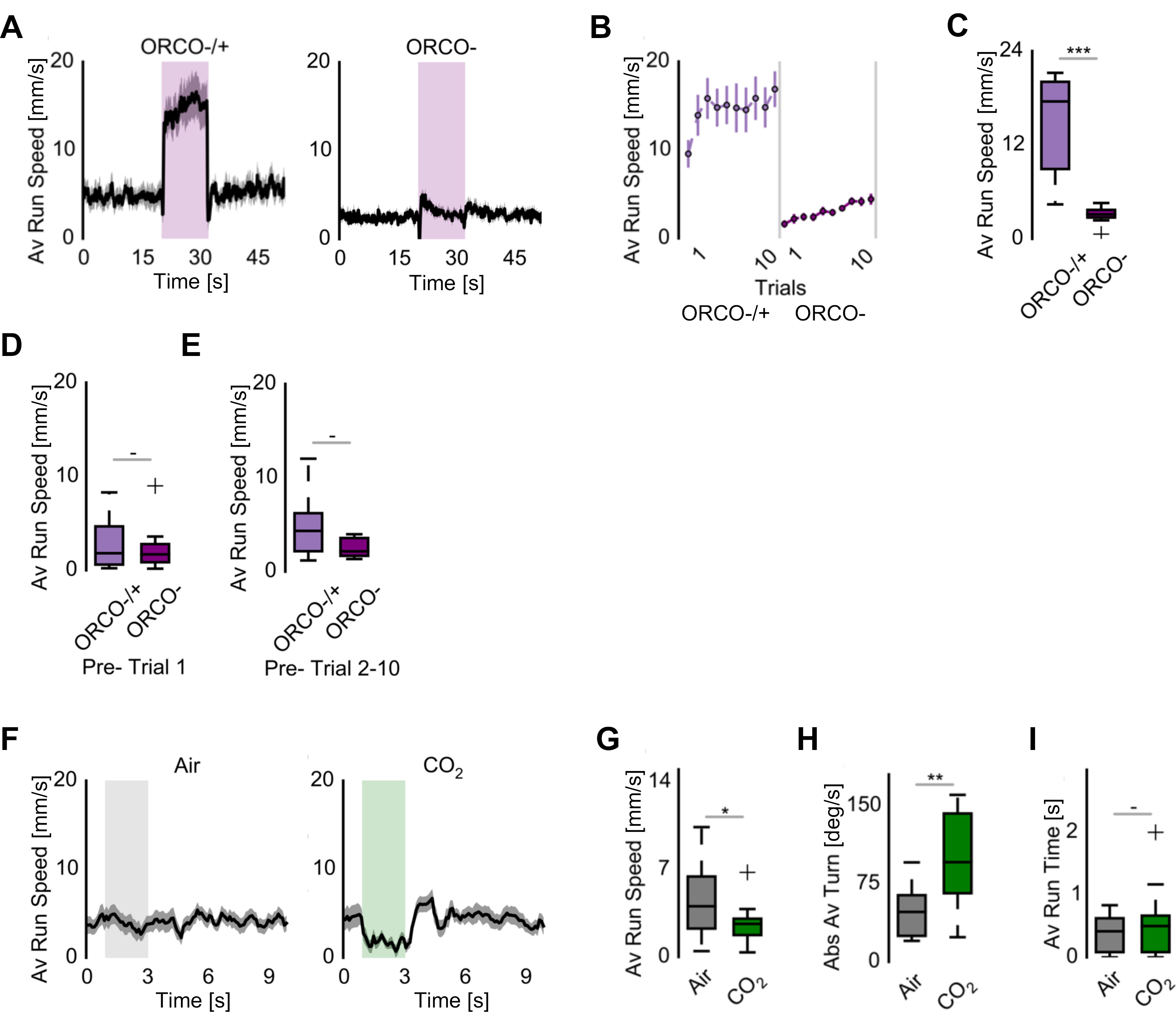
(A) Running behavior of food-deprived Orco null (Orco -) and heterozygous (Orco -/+) mutants. (B) Average running speed of Orco null and heterozygous mutants for vinegar over ten trials. (C) On average, Orco null mutants showed strongly reduced running after vinegar when compared to the heterozygous flies (n=10, paired T-test). (D) Running speeds recorded before odor exposure in the first trial for Orco null and heterozygous mutants. Naive flies were indistinguishable in locomotion and mobility (p= 0.725, n=10, T-test). (E) Running speeds recorded before odor exposure in Trials 2-10 for Orco null and heterozygous mutants. Orco -/+ flies showed a higher average in baseline speed, although this difference was not significant (p= 0.052, n=10, T-test) suggesting that odor detection is required for speed increase evolution. (F) Locomotion during aversive odor presentation. Flies were exposed to alternating CO_2_ and air plumes over six trials. (G) Flies slowed down at contact with aversive CO_2_. Average running speed under CO_2_ exposure was significantly lower than the speed observed during air exposure (n=10, paired T-test). (H) Flies executed escape turns under CO_2_ as they turned significantly more (n=10, paired T-test). (I) Initial running bout length did not differ between responses to air and CO_2_ (n=10, paired T-test). For all analyses, statistical notations are as follows: ‘ - ’ > 0.05, ’ * ‘ p < 0.05, ‘ ** ’ p < 0.01, ‘ *** ‘ p < 0.001.

**Fig. S2.**
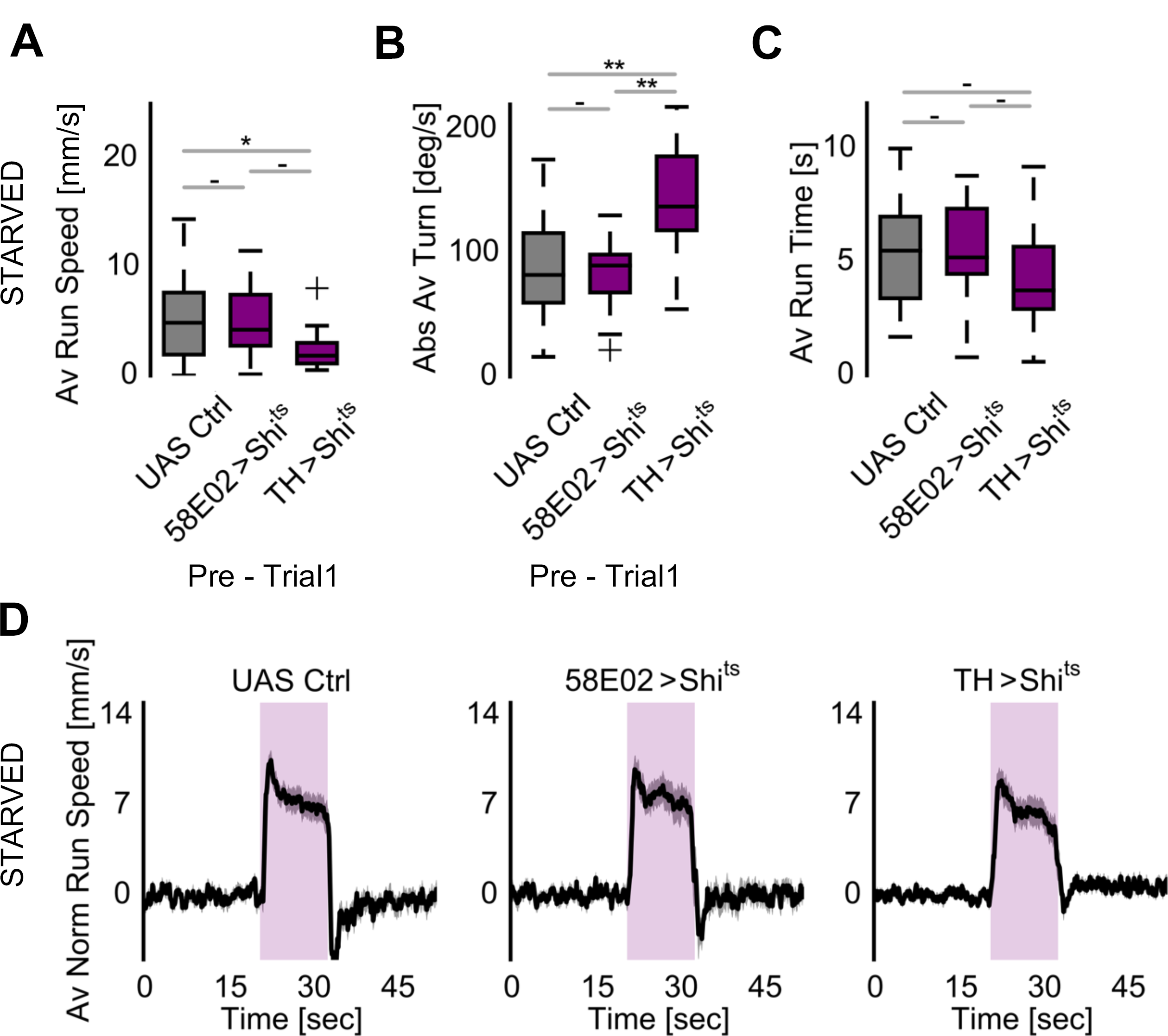
(A-B) Average running and absolute turning speeds for flies with thermogenetically silenced DANs prior to odor exposure in ten trials. Blocking PAM output (*58E02-Gal4>UAS-Shi^ts1^)* had no effect, while removing TH+/PPL1 DAN (*TH-Gal4>UAS-Shi^ts1^)* activity led to reduced running speed compared to only the control flies *(+>UAS-Shi^ts1^).* Meanwhile, without TH+/PPL1 DAN input, flies showed higher tendency to turn in either directions than PAM DAN block or the genetic controls (n=18/18/18). (C) Average running bout during odor stimulus for output silenced DANs. Blocking DAN output activity did not change the length of the non-interrupted running bout during odor stimulus (n=18/18/18). (D) Average normalized running speed upon transiently blocking DAN output. In the absence of PAM cluster *(58E02-Gal4>UAS-Shi^ts1^)* or TH+ positive *(TH-Gal4>UAS-Shi^ts1^)* dopaminergic neuron output, average running speeds during vinegar exposure were comparable to the genetic control after baseline normalization to the pre-stimulation phase suggesting that all groups accelerate equally in the presence of the odor stimulus. For all analyses, one-way ANOVA with Tukey’s HSD post hoc analysis was used: ‘ - ’ > 0.05, ’ * ‘ p < 0.05, ‘ ** ’ p < 0.01, ‘ *** ‘ p < 0.001.

**Fig. S3.1.**
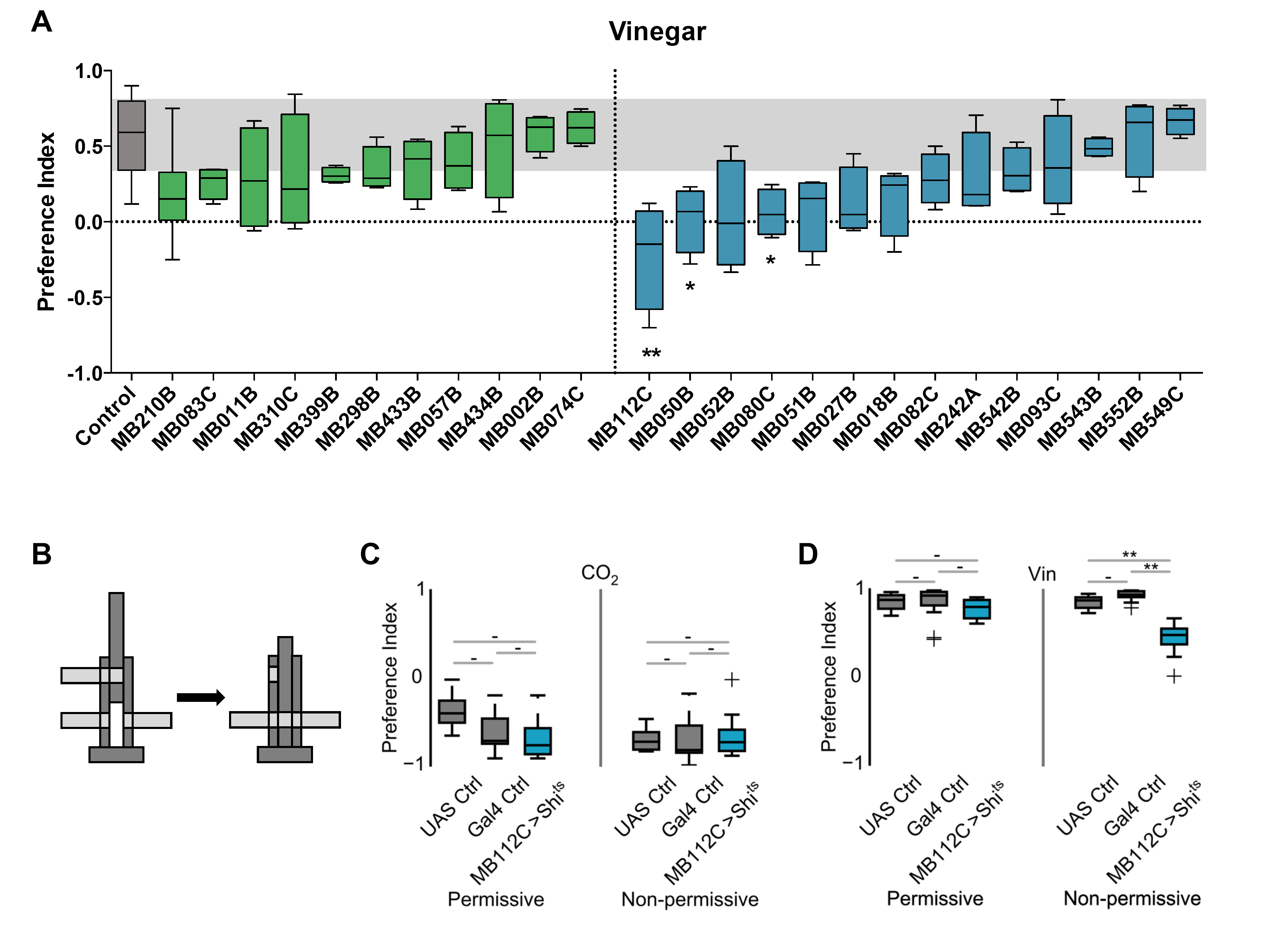
(A) T-maze neuronal silencing screen for mushroom body output neurons (MBONs) required for innate vinegar attraction. Several MB Split-Gal4 lines covering all MBON subsets were screened with the UAS-Shibire^ts1^ effector at the non-permissive temperature of 30°C (n=4-8 for experimental groups, controls were pooled to an n of 46-48, Kruskal-Wallis ANOVA and Dunn’s post-hoc test). Green: MBONs of horizontal lobe, blue: MBONs of vertical lobe. (B) Scheme of T-maze assay. Subsequent to loading of 40-60 flies into the T-maze, the elevator was lowered to the choice point. Flies were allowed to choose between tubes containing a test odor (vinegar 10% or 1% CO_2_) and a control (pressured air) for 1 minute. After 1 min, the elevator was moved back to the initial higher position and tubes were collected, to be counted later. Fly distribution was calculated as a preference index (# flies in odor - # flies in control) / ((# flies in odor + # flies in control). Permissive temperature (25°C) allows uninhibited synaptic transmission, whereas at non-permissive temperature (30°C), expression of Shibire^ts1^ blocks presynaptic neuron activity. (C-D) Preference indices for MB112C block in the T-maze for CO_2_ and vinegar. Manipulating MVP2 activity does not alter CO_2_ aversion in the T-maze (n=16/8/16 for permissive, n=16/8/16 for non-permissive). At permissive temperature, appetitive odor approach was not affected in all groups, however, blocking MVP2 in MB112C>Shi^ts1^ flies reduced attraction (n=16/8/16 for permissive, n=16/8/16 for non-permissive, one-way ANOVA with Tukey’s HSD post hoc analysis). For all analyses, statistical notations are as follows: ‘ - ’ > 0.05, ’ * ‘ p < 0.05, ‘ ** ’ p < 0.01, ‘ *** ‘ p < 0.001.

**Fig. S3.2.**
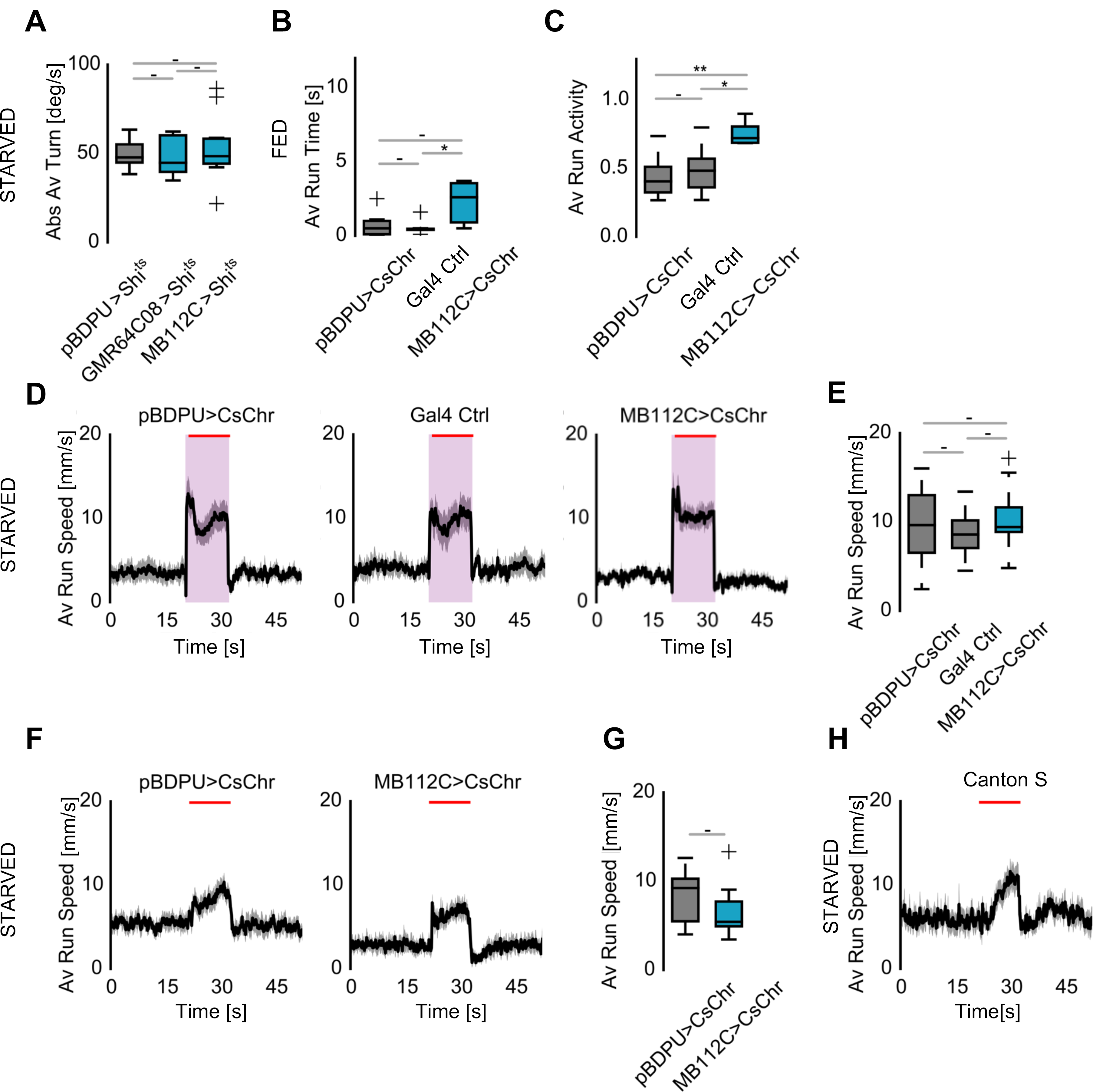
(A) Average absolute turning speeds for flies with synaptically-blocked MVP2 and mushroom body γ Kenyon cells (n=10/10/10). (B) Average running bout times for MVP2 activation in fed flies (n=7/7/7). Activation of MVP2 (*MB112C>UAS-CsChrimson)* increased the first running bout length only compared to the Gal4 control *(MB112-Gal4>+)*(Tukey HSD q-statistics score between *MB112C>UAS-CsChrimson* and UAS-Gal4 control was 3.525 (p < .05 (q-critical[3, 17] = 3.627)). (C) Average running activity during odor stimulation period for MVP2 activation in fed flies. When MVP2 was activated *(MB112C>UAS-CsChrimson),* flies were more active in running in comparison to controls (n=7/6/7, controls: *pBDP-Gal4U>UAS-CsChrimson* and *MB112-Gal4>+).* (D,E) Average running speeds in starved flies when MB112C was activated via optogenetics *(MB112C>UAS-CsChrimson).* Activation of MVP2 did not further increase odor tracking behavior of starved flies (n=10/10/10, controls: *pBDP-Gal4U>UAS-CsChrimson* and *MB112-Gal4>+).* (F,G) Running speed averages for the light only condition of *MB112C>UAS-CsChrimson* and empty-Gal4 control *(pBDP-Gal4U>UAS-CsChrimson)* animals. MVP2 activation in starved flies did not induce locomotion or tracking compared to the control flies in the absence of odor stimulation (n=10/10, T-test). (H) Wild-type control flies show a mild increase in motility and locomotion under 617nm, 30W/mm2 light delivery (n=10). The flies are presumably attracted to the light in the absence of CsChrimson expression. For all analyses, one-way ANOVA with Tukey’s HSD post hoc analysis was used: ‘ - ’ > 0.05, ’ * ‘ p < 0.05, ‘ ** ’ p < 0.01, ‘ *** ‘ p < 0.001.

**Fig. S3.3.**
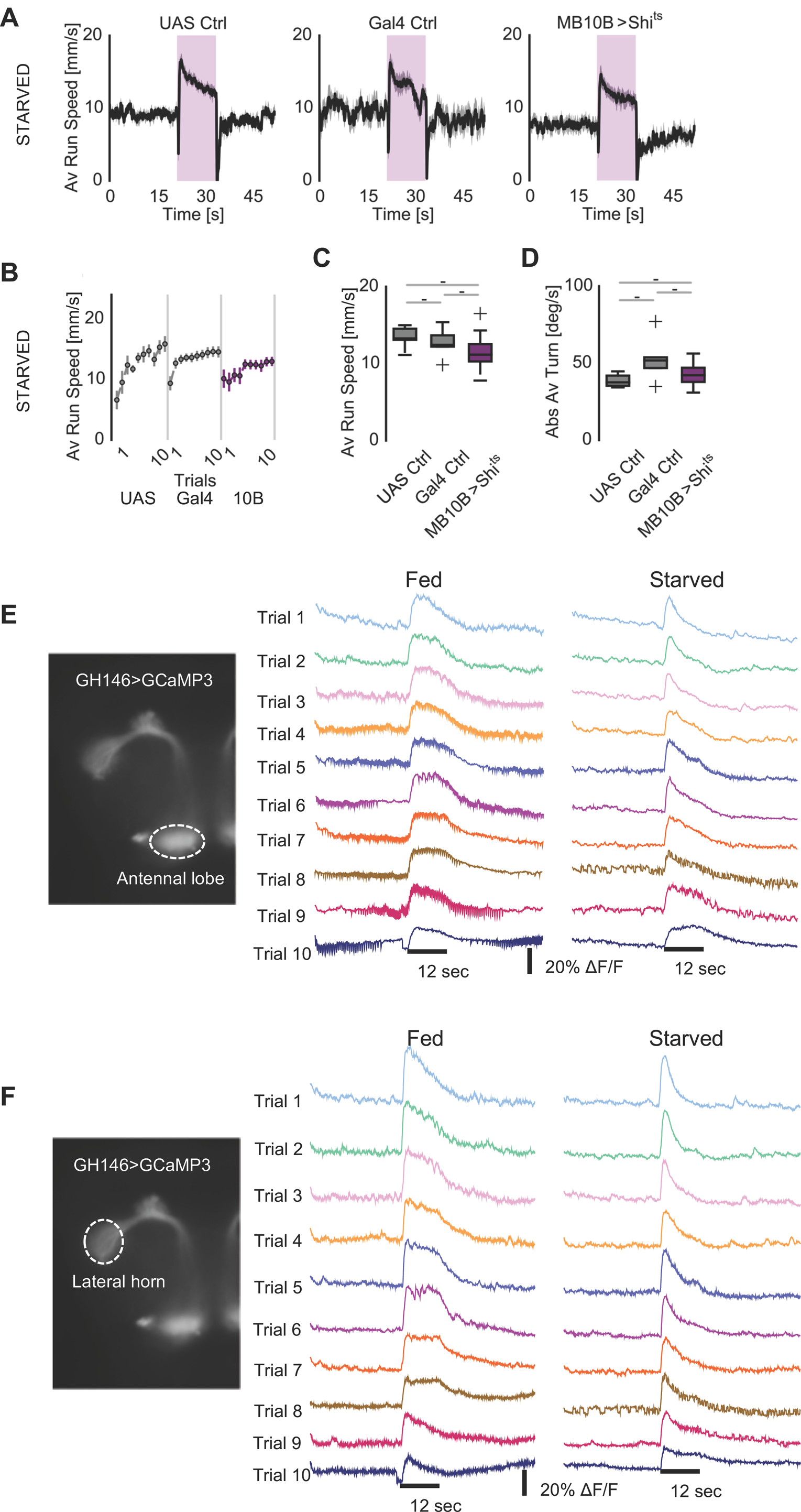
(A-D) Inactivation of all Kenyon cell synaptic output *(MB10B-Gal4>UAS-shi^ts1^)* did not affect odor tracking behavior of 24 h starved flies compared to genetic controls (n=5/5/7, controls: UAS-Ctrl *+>UAS-shi^ts1^*, Gal4-Ctrl *MB10B-Gal4>+,* one-way ANOVA with Tukey’s HSD post hoc analysis: ‘ - ’ > 0.05, ’ * ‘ p < 0.05, ‘ ** ’ p < 0.01, ‘ *** ‘ p < 0.001.). (E) *In vivo* calcium imaging of the AL region in starved and fed flies. Traces show the responses of PNs *(GH146-Gal4>UAS-GCaMP3)* upon repeated stimulation with vinegar (12 s/trial). Note that the response does not decrease or increase significantly over repeated stimulation trials, and it is not different between starved and fed animals. (F) *In vivo* calcium imaging of the lateral horn (LH) region in starved and fed flies. Traces show the responses of PNs *(GH146-Gal4>UAS-GCaMP3)* upon repeated stimulation with vinegar (12 s/trial). Responses did not change between trials or feeding state.

**Fig. S4.**
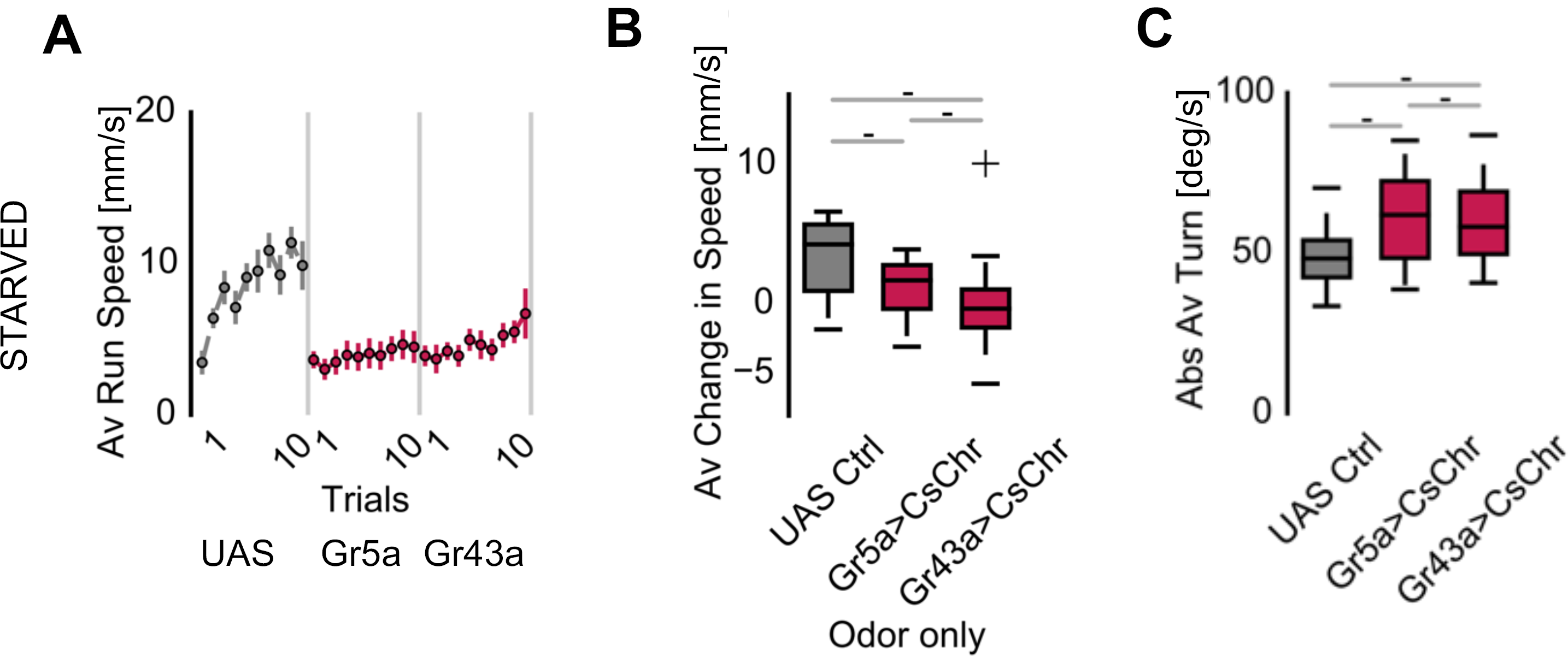
(A) Evolution of average running speeds over ten trials for heterozygous control (+>UAS-*CsChrimson)* and *Gr>UAS-CsChrimson* flies (n=10/10/10). (B) Average normalized running speed during 2 s odor only phase for control and *Gr>UAS-CsChrimson* flies. The average change in speed was calculated by normalization of the average speeds of trial 2-10 by subtracting the average speed of trial 1 (n=10/10/10, one-way ANOVA with Tukey’s HSD post hoc analysis). Odor tracking speed does not significantly increase upon pairing of the odor with gustatory neuron activation between first odor exposure in naive animals as compared to animals that experienced pairing. (C) Average absolute turning speeds during Gr neuron activation (n=10/10/10). For all analyses, statistical notations are as follows: ‘ - ’ > 0.05, ’ * ‘ p < 0.05, ‘ ** ’ p < 0.01, ‘ *** ‘ p < 0.001.

**Fig. S5.1.**
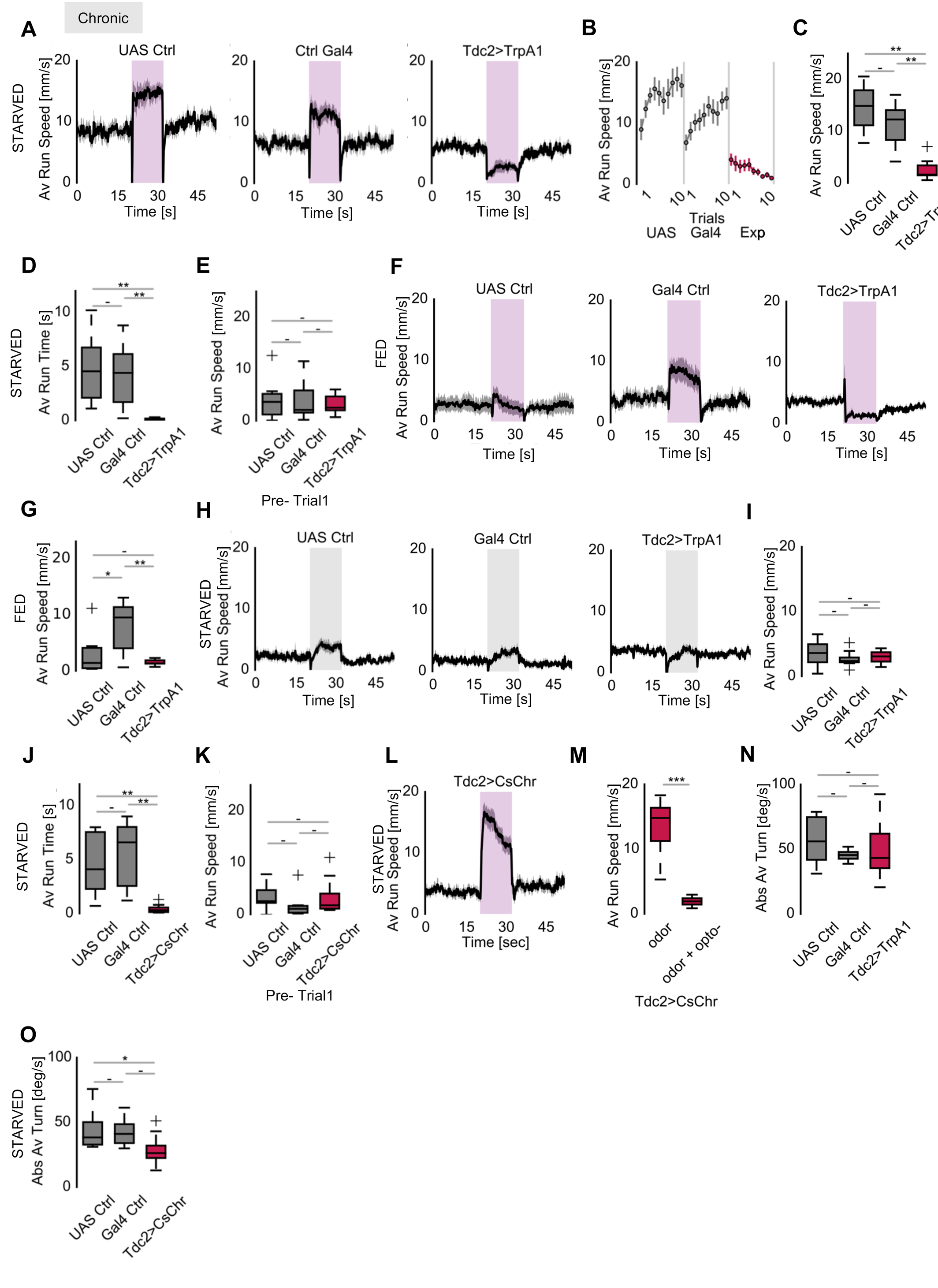
(A) Thermogenetic activation of octopaminergic neurons during odor tracking. *Tdc2-Gal4* drives the heat-inducible cation channel dTrpA1 in octopaminergic neurons. Compared to the heterozygous genetic controls (UAS Ctrl: *+>UAS-dTrpA1*, Gal4 Ctrl: *Tdc2-Gal4>+),* depolarizing octopaminergic neurons led to abrupt stopping or slowing down at odor exposure. This was not the case in the controls. (B) Evolution of average running speed for *Tdc2>UAS-dTrpA1* flies and controls during odor exposure over trials. (C) Average running speeds for *Tdc2>UAS-dTrpA1* flies during odor exposure. On average, the running speed during active odor tracking was significantly reduced due to octopaminergic neuron activation (n=10/10/10, one-way ANOVA with Tukey’s HSD post hoc analysis). (D) Odor stimulation resulted in immediate stop when octopaminergic neurons were artificially activated with TrpA1 *(Tdc2>UAS-TrpA1)* (n=10/10/10). (E) Average running speeds for flies prior to odor exposure during chronic activation of Tdc2+ neurons (n=10/10/10). Note that odor stimulus-independent running speeds do not differ between test and control groups. (F-G) Thermogenetic chronic activation of octopaminergic neurons in fed flies (n=9/8/8). The Tdc2-Gal4 control frequently showed some odor-induced tracking even in fed flies. (H-I) Average running speeds for the air only response in chronic activation of Tdc2+ neurons (n=10/10/10). No significant decrease was observed in the average speed during air stimulation (I). (J) Running times for acute Tdc2 activation experiments during odor stimulation. Tdc2 activated flies tracked the odor for significantly shorter times (n=10/10/10). (K) Average running speeds for flies prior to first odor exposure of *Tdc2>UAS-CsChrimson* flies (n=10/10/10). (L) Average running speeds of *Tdc2>UAS-CsChrimson* flies to odor only stimulation without any optogenetic stimulation (n=10). As seen before, vinegar induces strong tracking in starved flies. (M) Comparison of running speeds of *Tdc2>UAS-CsChrimson* flies in odor only and during odor + optogenetic stimulation (Combined data from Figure 4D, n=10/10, T-test). (N) Average absolute turning speeds during vinegar response in flies with chronically activated Tdc2+ neurons (n=10/10/10). (O) Average absolute turning speeds for flies during vinegar response for *Tdc2>UAS-CsChrimson* and control (UAS Ctrl: *+>UAS-CsChrimson,* Gal4 Ctrl: *Tdc2-Gal4>+)* flies (n=10/10/10). Turning behavior was reduced in flies when octopaminergic neurons were activated (Tukey HSD q-statistics score between *Tdc2>CsChrimson* and Gal4 control was 3.351 (p < .05 (q-critical[3, 27] = 3.50)). For all analyses, except Fig. S5.1M, one-way ANOVA with Tukey’s HSD post hoc analysis was used. : ‘ - ’ > 0.05, ’ * ‘ p < 0.05, ‘ ** ’ p < 0.01, ‘ *** ‘ p < 0.001.

**Fig. S5.2.**
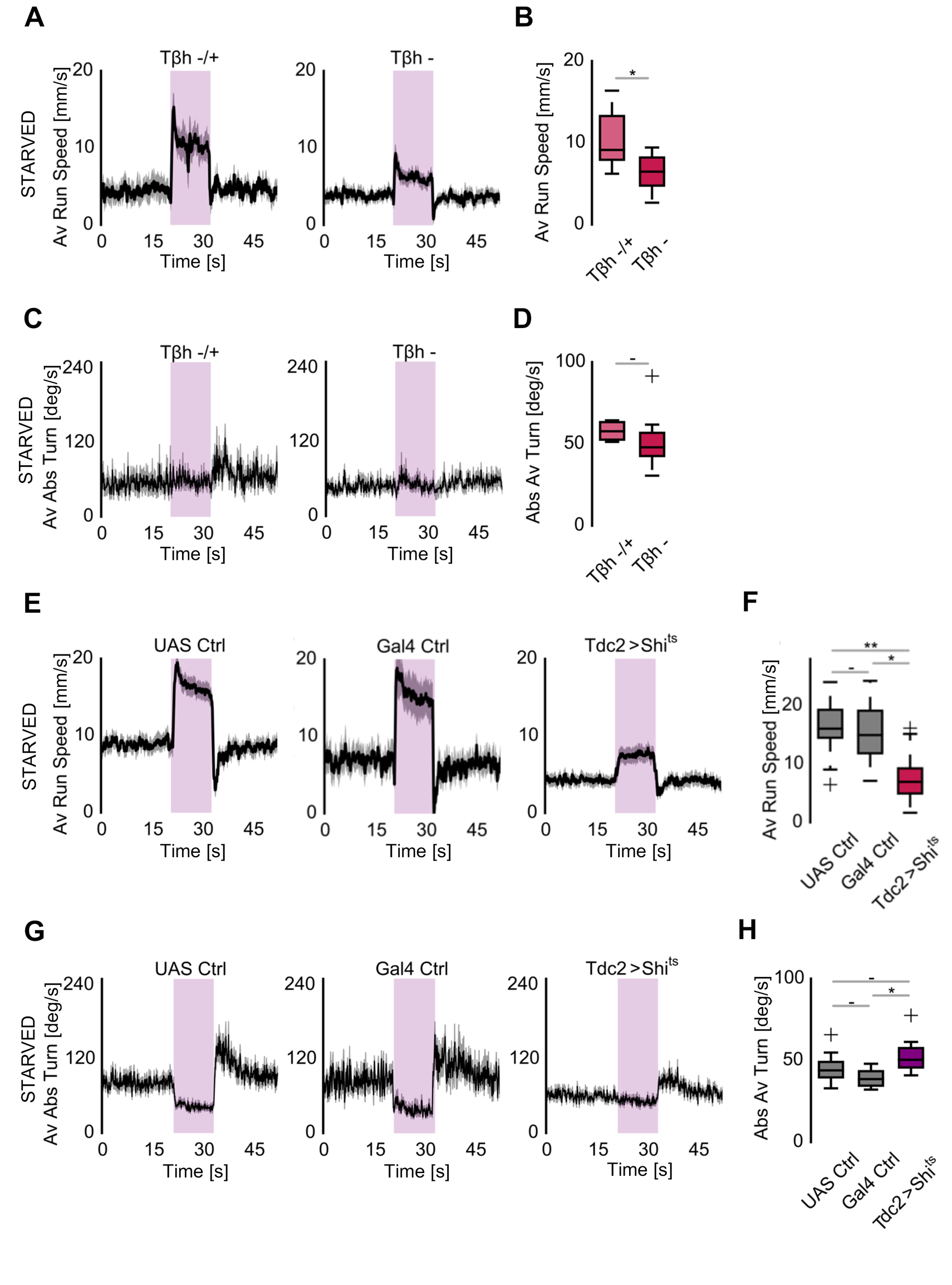
(A) Average running speeds of TβH heterozygous (-/+) and null mutants (-). TβH mutants cannot produce octopamine (OA) from tyramine. Permanent loss of OA leads to an overall reduced running speed consistent with a role of systemic OA in arousal. (B) Average running speeds of TβH heterozygous and null mutants in ten trials (n=8/5, T-test). (C) Average absolute turning speeds of TβH heterozygous (-/+) and null mutants (-). (D) Average absolute turning speeds of TβH heterozygous and null mutants in ten trials (n=8/5, T-test). (E) Average running speeds for *Tdc2>Shi^ts1^* and genetic control flies with blocked synaptic transmission from OA synapses. This phenotype is again consistent with a systemic role of OA in arousal and suggests that multiple OA neurons with partly antagonistic functions are contributing to the phenotypes observed (UAS control same as in Fig. 2A: *+/UAS-shi^ts1^,* Gal4 control: *Tdc2-Gal4>+).* (F) Blocking Tdc2+ synaptic output reduced odor approach speed (n=18/9/18, one-way ANOVA with Tukey’s HSD post hoc analysis). (G-H) Average absolute turning speeds for blocking of octopaminergic neuron output. Blocking of Tdc2+ neurons did not change turning behavior (UAS control same as in Fig. 2D: *+/UAS-shi^ts1^,* Gal4 control: *Tdc2-Gal4>+,* n=18/9/18, one-way ANOVA with Tukey’s HSD post hoc analysis). For all analyses, statistical notations are as follows: ‘ - ’ > 0.05, ’ * ‘ p < 0.05, ‘ ** ’ p < 0.01, ‘ *** ‘ p < 0.001.

**Fig. S6.1.**
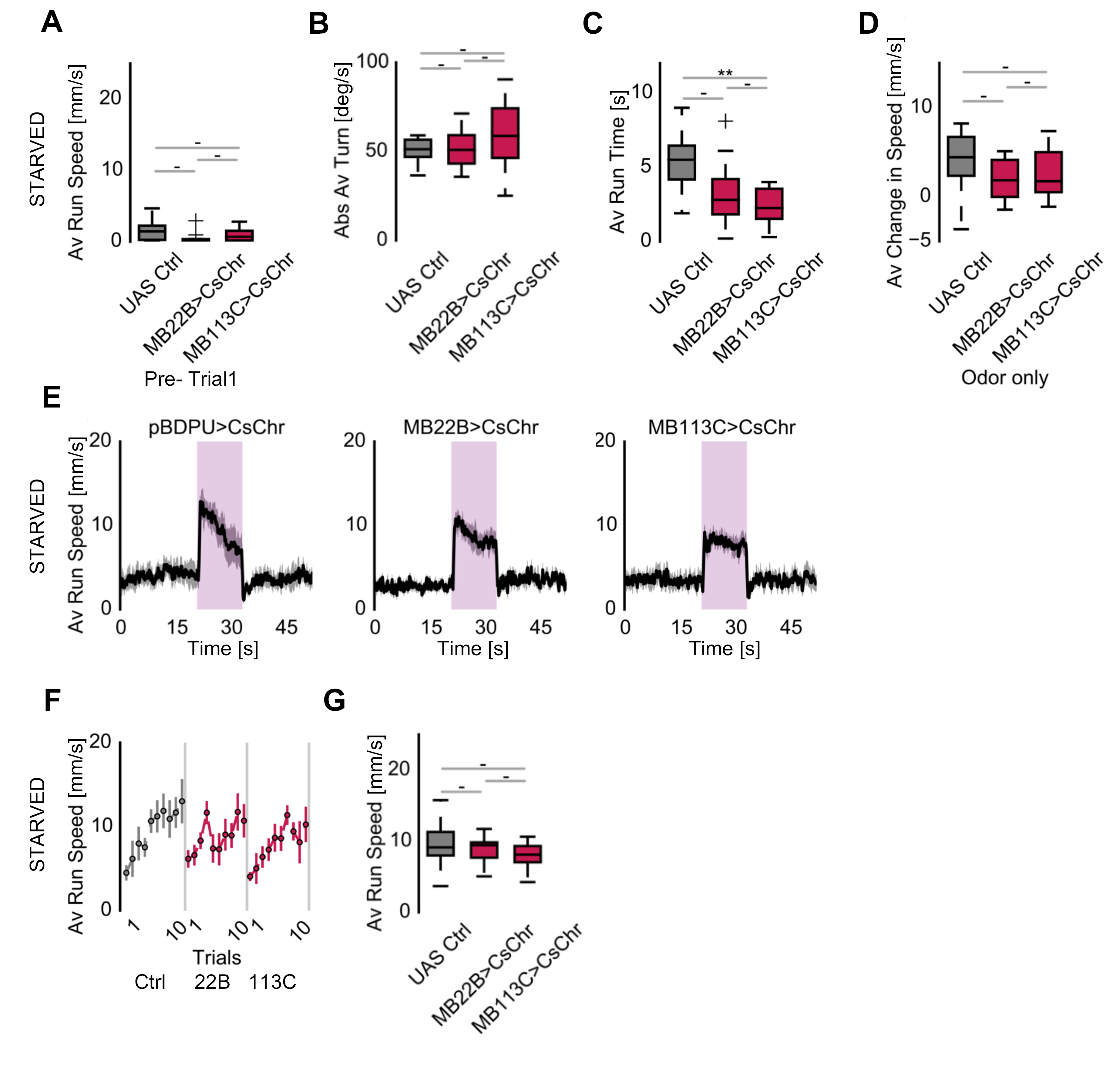
(A) Average running speeds for VPM lines (*MB22B>UAS-CsChrimson* and *MB113C>UAS-CsChrimson)* and the control *(pBDP-Gal4U>UAS-CsChrimson)* before odor exposure. These lines did not exhibit motor defects. (n=10/10/10). (B) Average absolute turning speeds for VPM lines during concurrent odor and optogenetic presentation. Activating VPM neurons did not alter turning behavior (n=10/10/10). (C) Average running bout times for VPM neurons. Running bout length was reduced with activation of MB113+ neurons. (n=10/10/10, Tukey HSD q-statistics score between *MB22B>UAS-CsChrimson* and Gal4 control was 3.459 (p < .05 (q-critical [3, 27] = 3.50)). (D) Average normalized change of running speed during only odor exposure. To test the whether activating VPMs artificially induce odor learning, running speeds recorded in odor only time window of the later trials (Trial 2-10) were normalized to the first trial odor only behavior. *MB22B>UAS-CsChrimson* or *MB113C>UAS-CsChrimson* flies performed in a statistically comparable way to the control group (n=10/10/10). (E-G) Average running speeds of *MB22B>UAS-CsChrimson* and *MB113C>UAS-CsChrimson* flies under vinegar in the absence of optogenetic stimulation. These flies were indistinguishable from the control *pBDP-Gal4U>CsChr* (n=6/6/6). For all analyses, one-way ANOVA with Tukey’s HSD post hoc analysis was used. : ‘ - ’ > 0.05, ’ * ‘ p < 0.05, ‘ ** ’ p < 0.01, ‘ *** ‘ p < 0.001).

**Fig. S6.2.**
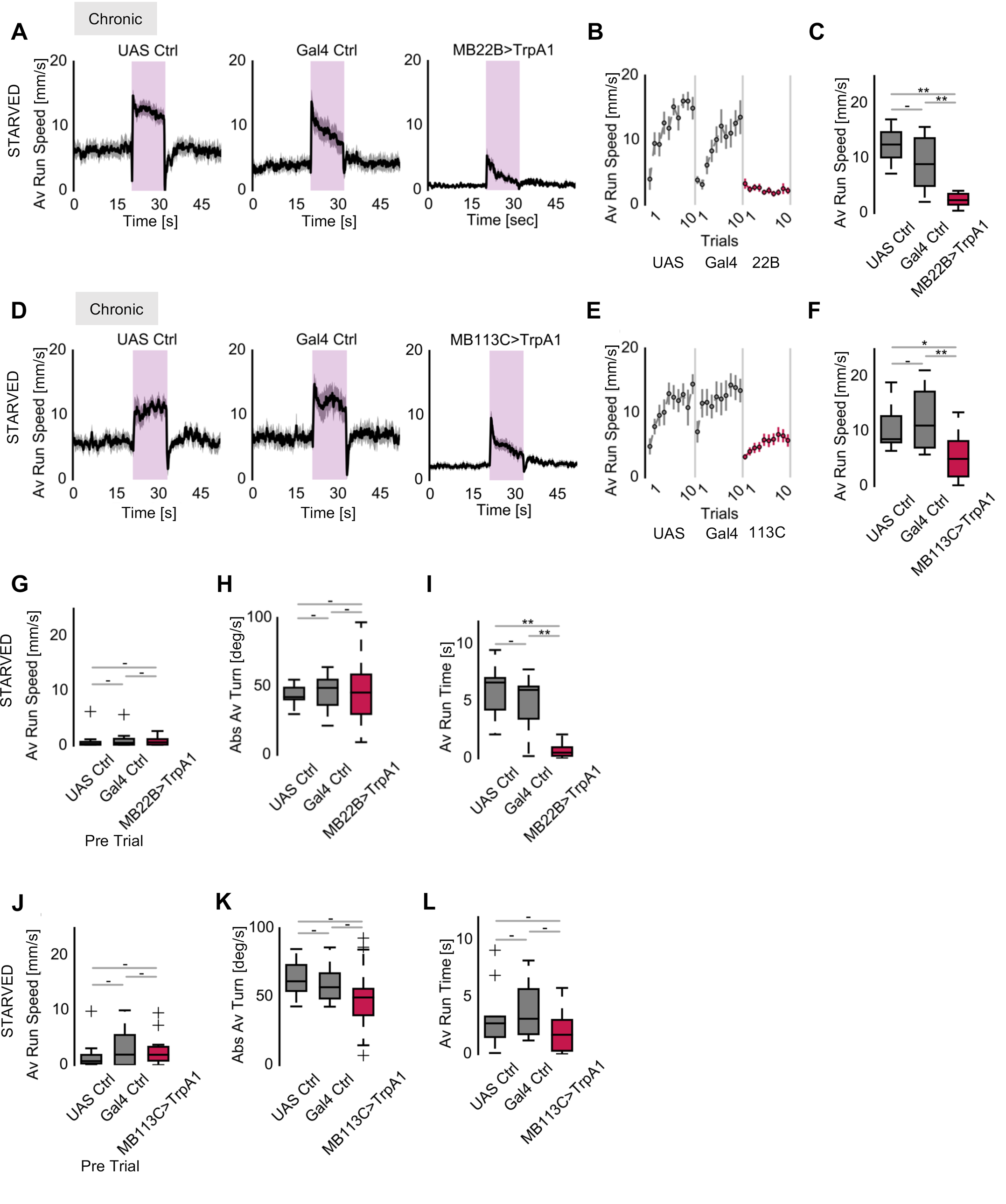
(A-L) Thermogenetic chronic activation of VPM neurons. At 30°C, *MB22B>UAS-dTrpA1 activated* VPM3 and VPM4, whereas *MB113C>UAS-dTrpA1* targets activation of only VPM4 neurons chronically. The genetic controls are UAS Ctrl: *+>UAS-dTrpA1* and Gal4 Ctrl: *MB22B>+* and *MB113C>+* respectively. Activation of VPM neurons impeded vinegar tracking in hungry flies (A-F) (n=10/10/10 for A-C and n=10/9/29 for D-F). (H) Average running speeds prior to first odor encounter for chronic VPM activation. Expression of dTrpA1 in VPM neurons did not cause motor defects (G, J) (n=10/10/10 for G, n=10/9/29 for J). Average absolute turning speeds for chronic activation of VPMs. Activating VPMs did not alter turning response (H, K) (n=10/10/10 for H, n=10/9/29 for K). Average running bout times for VPMs chronic activation during odor stimulation. Combined activation of VPM3 and VPM4 resulted in earlier stops and shorter running bouts (I, L) (n=10/10/10 for I, n=10/9/29 for L). For all analyses, one-way ANOVA with Tukey’s HSD post hoc analysis was used. : ‘ - ’ > 0.05, ’ * ‘ p < 0.05, ‘ ** ’ p < 0.01, ‘ *** ‘ p < 0.001.

**Fig. S6.3.**
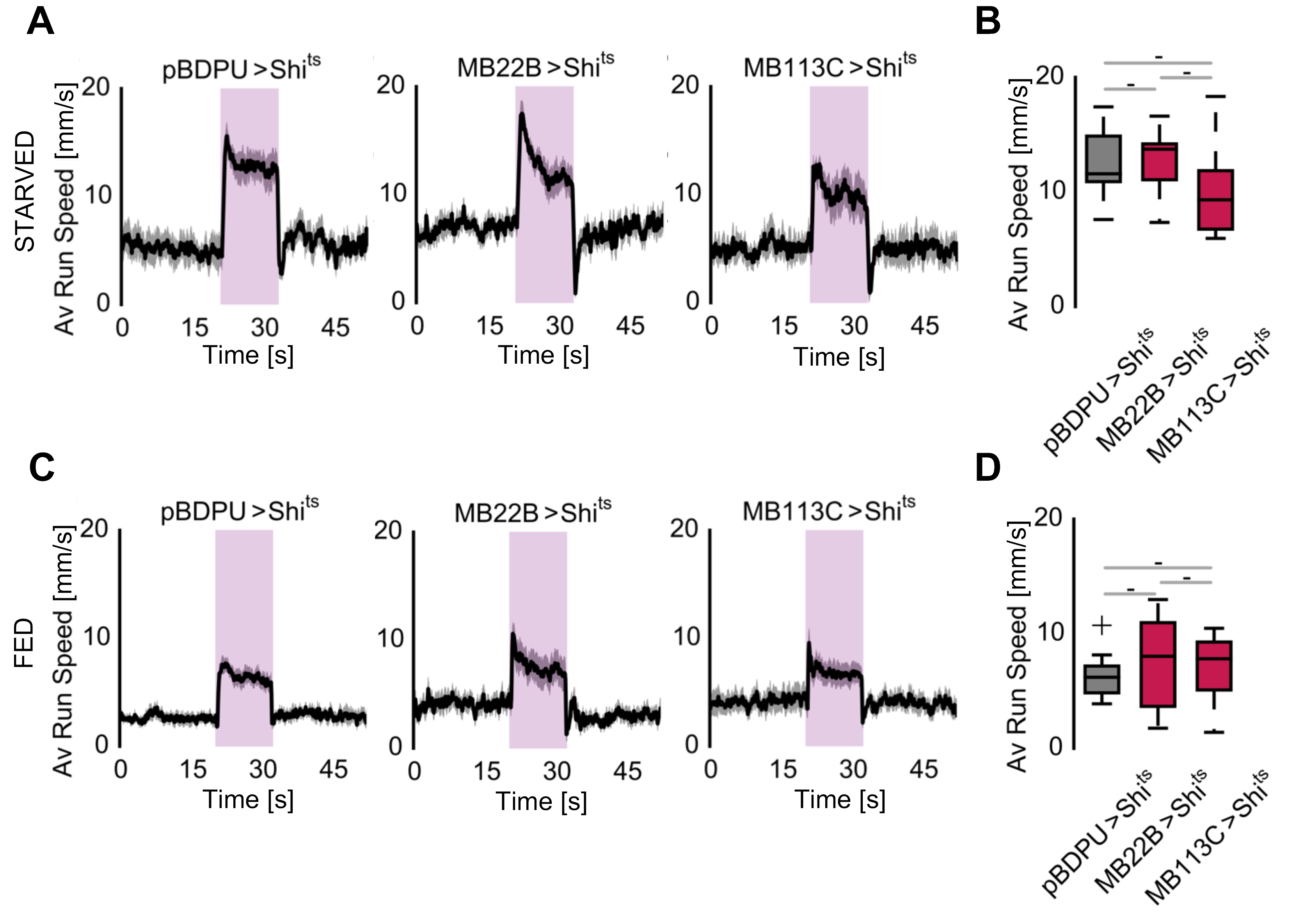
(A-D) In fed and starved flies, running speed in average during synaptic activity blocking of VPM neurons *(MB22B>UAS-CsChrimson* and *MB113C>UAS-CsChrimson)* and controls *(pBDP-Gal4U>UAS-CsChrimson).* Perseverant running speed under vinegar did not change in flies with VPM neuron output blocked. (n=7/8/8 for Fig. S6.3A-B, n=10/10/10 for Fig. S6.3C-D). For all analyses, one-way ANOVA with Tukey’s HSD post hoc analysis was used. : ‘ - ’ > 0.05, ’ * ‘ p < 0.05, ‘ ** ’ p < 0.01, ‘ *** ‘ p < 0.001.

**Fig. S7.1.**
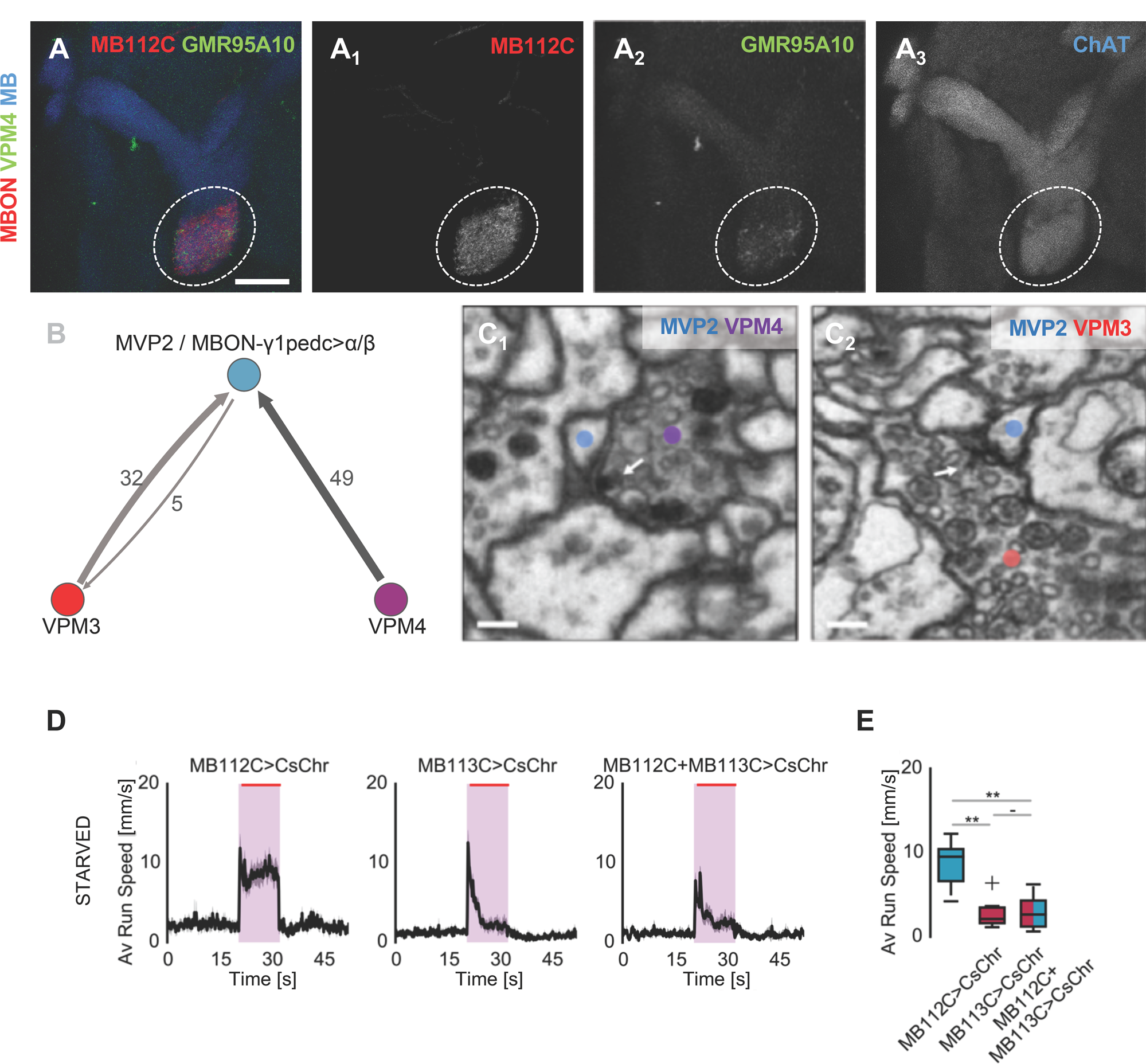
(A, A_1-3_) Double stainings suggest that MVP2 and VPM4 neurons could be synaptically connected. In the double labeling experiment, MVP2 was visualized by *MB112C-Gal4>UAS-mCD8-RFP* (red) and VPM4 neurons were labeled by *GMR95A10-LexA>LexAop2-mCD8-GFP* (green). (A) Merged image displaying MVP2 and VPM4 neurons. MB lobes are stained with anti-Chat (blue) (A_1-3_, single channels; 10μm scale bar. Note that VPM4 neurons innervate the dendritic region of MVP2 at the level of the MB γ1 lobe region. (B) Wiring diagram of VPM3, VPM4 and MVP2 connectivity. Both VPM3 and VPM4 target MVP2 dendrites in the γ1 compartment. Only VPM3 is reciprocally connected with MVP2. Numbers represent synapse counts. (C1,C2) Representative synapses (arrow) between VPM3 (red), VPM4 (purple) and MVP2 (blue) show both clear core vesicles around the active zone as well as large dense core vesicles in close vicinity. 100nm scale bar (D,E) Average running speeds in starved flies when MB112C and VPM4 neurons were activated via optogenetics simultaneously (*MB112C-Gal4; MB113C-Gal4>UAS-CsChrimson).* Activation of MVP2 did not further increase appetitive odor response and tracking in starved animals. VPM4 and MVP2 co-activation in starved flies. Note that VPM4 activation completely suppresses odor tracking in the starved animal as seen also in the fed animal suggesting that VPM4 inhibits odor tracking regardless of starvation state (n=7/7/7). For all behavioral analyses, one-way ANOVA with Tukey’s HSD post hoc analysis was used. : ‘ - ’ > 0.05, ’ * ‘ p < 0.05, ‘ ** ’ p < 0.01, ‘ *** ‘ p < 0.001.

**Fig. S7.2.**
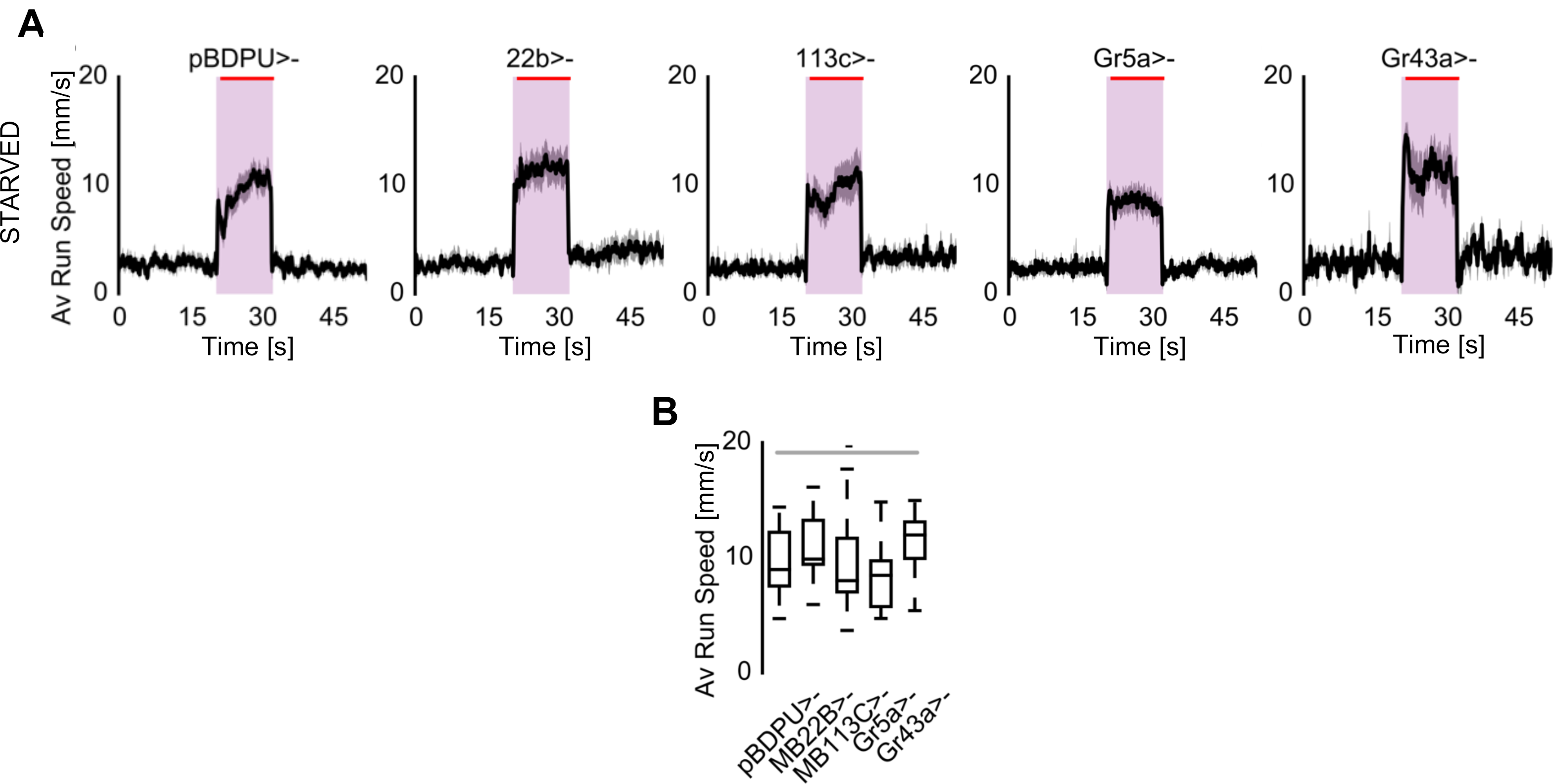
(A-B) Gal4 controls for split-Gal4 and gustatory receptor neuron Gal4-lines during simultaneous vinegar and optogenetic application (n=10/10/10/10/4). No heterozygous Gal4 only control behaved different from empty-Gal4 *(pBDP-Gal4U>UAS-CsChrimson)* control. For all behavioral analyses, one-way ANOVA with Tukey’s HSD post hoc analysis was used. : ‘ - ’ > 0.05, ’ * ‘ p < 0.05, ‘ ** ’ p < 0.01, ‘ *** ‘ p < 0.001.

